# Genetic, intrinsic, and environmental determinants of innate immune cytokine responses in healthy four-year-old children

**DOI:** 10.64898/2026.04.30.722087

**Authors:** Rutger J. Röring, Luba Sominsky, Katherine Lange, Aaron L. Weinman, Jim Buttery, Rachel J. Morgan, Gabrielle P.D. MacKechnie, Kavindi Gamage, Katherine Drummond, Peter Sly, Fiona Collier, Anne-Louise Ponsonby, Markus Juonala, Deborah A. Lawlor, Petter Brodin, Mihai G. Netea, Niels P. Riksen, Mimi L.K. Tang, Boris Novakovic, Richard Saffery, Peter Vuillermin, Toby Mansell, David P. Burgner, the Barwon Infant Study investigator group

**Affiliations:** Murdoch Children’s Research Institute, Parkville, Victoria, Australia; Institute for Mental and Physical Health and Clinical Translation (IMPACT), School of Medicine, Deakin University, Geelong, Victoria, Australia; Barwon Health, Geelong, Victoria, Australia; Department of Paediatrics, The University of Melbourne, Melbourne, Victoria, Australia; Epidemiology Informatics Group, Murdoch Children’s Research Institute, Parkville, Victoria, Australia; The Florey Institute of Neuroscience and Mental Health, University of Melbourne, Parkville, Victoria, Australia; Child Health Research Centre, The University of Queensland, Brisbane, Queensland, Australia; Department of Internal Medicine, University of Turku, Turku, Finland and Division of Medicine, Turku University hospital, Turku, Finland; MRC Integrative Epidemiology Unit at the University of Bristol, Oakfield House, Oakfield Grove, Bristol, BS8 2BN, UK; Population Health Sciences, Bristol Medical School, University of Bristol, Oakfield House, Oakfield Grove, Bristol, BS8 2BN, UK; Unit for Clinical Pediatrics, Department of Women’s and Children’s Health, Karolinska Institutet, 17165 Solna, Sweden; Department of Immunology and Inflammation, Imperial College London, W12 0NN London, UK; Medical Research Council Laboratory of Medical Sciences (LMS), Imperial College Hammersmith Campus, London, UK; Dept. of Internal Medicine, Radboud university medical center, Nijmegen, the Netherlands; Department of Immunology and Metabolism, Life & Medical Sciences Institute, University of Bonn, Bonn, Germany; Department of Allergy and Immunology, Royal Children’s Hospital, Melbourne, Victoria, Australia

## Abstract

Innate immune responses are crucial for host defence but vary markedly between individuals. Although determinants of this variation are well characterised in adults, data from healthy children remain scarce. We therefore profiled whole-blood cytokine responses to innate immune stimulation in 286 children aged approximately four years and examined genetic, host-intrinsic, and environmental correlates of response. Cytokine responses showed marked inter-individual heterogeneity and stimulus-specific patterns. The top 50 genetic variants explained a substantial proportion (∼20-45%) of this variance across many stimulus conditions, including a biologically coherent association of the STING locus with cGAMP-induced cytokine production. In contrast, sex, age, adiposity, and perinatal variables showed limited or modest associations. Systemic inflammatory biomarkers of systemic inflammation (hsCRP, glycoprotein acetyls, granulocyte-to-lymphocyte ratio) were strongly positively associated with cytokine responses. Finally, seasonal population-level viral infection burden was positively associated with antiviral and inflammatory cytokine responses. Collectively, these findings advance our understanding of variation in early-life whole-blood cytokine responses, underscoring this developmental period as a critical window for understanding immune development trajectories relevant to long-term health.

## Introduction

Innate immune responses, including cytokine production, are central to immediate protection from infection and injury^1^. They also contribute to the pathophysiology of chronic inflammatory diseases including atherosclerosis^2^. Susceptibility to these infectious and noncommunicable diseases is partly determined by marked variation (including dysregulation) in innate immune responses between individuals, across the life course^1,3–10^. In healthy adults, innate immune cytokine responses to *ex-vivo* stimulation are shaped by both genetic variation and environmental exposures such as diet and infection^6–8,11,12^. The heritability of these innate responses in adults varies by stimulus and cytokine, is generally higher than for adaptive responses^7^, and proportionally declines with age^13^.

In early life, notwithstanding rapid maturation and development, immune responses differ markedly from those in adults, with a relative reliance on innate immune responses rather than adaptive responses^3,14–16^. Despite their importance in childhood and for lifetime disease risk, the determinants of innate immune cytokine responses in childhood are poorly understood. Few studies have characterised innate immune responses in healthy children - existing paediatric data are largely derived from those with immune-related conditions (such as allergy or asthma)^17,18^ or from studies of the effects of specific immune-modulating agents (such as Bacillus Calmette–Guérin vaccine^19–21^). This limits understanding of how inter-individual variation in innate responses is established during early life.

To address this gap, we profiled innate immune cytokine responses in whole blood samples from a large sample of 4-year-olds in a deeply phenotyped longitudinal cohort with detailed participant information and repeated biological samples^22^. We investigated the contribution of genetic (quantitative trait locus [QTL]-specific and overall heritability), non-genetic host (e.g. birth factors, sex, anthropometry) and environmental (e.g. seasonal variation in population-level incidence of respiratory viral infection) determinants of innate cytokine responses at four years of age, and examined how of these responses relate to systemic inflammation and leukocyte composition.

## Results

### Innate immune stimulation reveals stimulus-specific and highly variable cytokine responses in early childhood

To investigate cytokine responses in four-year-old children, we used samples collected as part of the Barwon Infant Study (BIS), a population-derived Australian cohort of mainly (approximately 90%) European descent^22^. Briefly, fresh peripheral blood samples from 286 children were incubated for 24 hours with a panel of 8 pathogen mimetics or appropriate controls, and 13 innate immune cytokines were quantified (Figure 1A). Details regarding the stimuli and cytokines are provided in the Methods section (see also Table 1).

**Figure 1:**
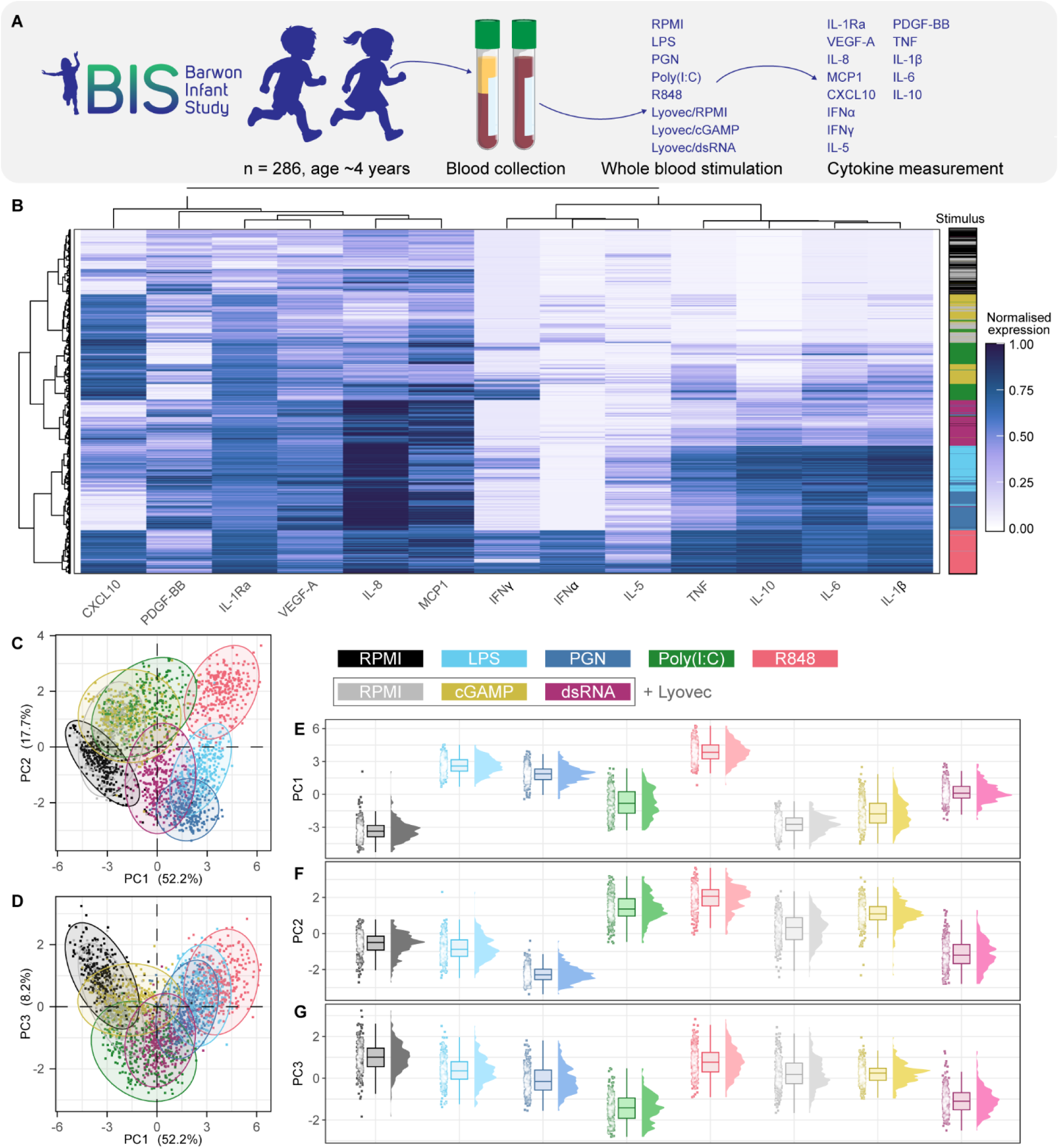
Describing innate immune cytokine responses in pre-school children. (A) Schematic overview of the study procedure. (B) Intensity-normalised (‘min-max standardised’) heatmap of all cytokine responses. Rows: individual samples (1 sample per stimulus per child). Columns: individual cytokines. Clustering was done using Ward’s method with Euclidian distance. (C-D) Principal component analysis of all samples, coloured by stimulus. (E-G) Separate visualisation of principal components 1, 2, and 3, coloured by stimulus. Points represent individual samples; summary boxplot and distribution are additionally provided. All data were adjusted for time in freezer and exact incubation time of the assay. For each stimulus, dots represent individual samples with the boxplot and Boxplots are in the style of Tukey with the median and IQR between the 25^th^ and 75^th^ percentile.

**Table 1:** Details of stimuli for whole-blood stimulations.

| Stimulus | Concentration | Manufacturer | Notes |
| --- | --- | --- | --- |
| RPMI | - | Invitrogen |  |
| LPS | 100 ng/mL | Invivogen | *E. coli* O111:B4 |
| PGN | 10 µg/mL | Invivogen |  |
| R848 | 10 µM (~3.5 µg/mL) | Invivogen |  |
| Poly(I:C) | 100 µg/mL | Invivogen |  |
| Lyovec | 10% (v/v) | Invivogen |  |
| Lyovec/cGAMP | 10 µg/mL | Invivogen |  |
| Lyovec/dsRNA | 1 ng/mL | invivogen |  |

The cytokine production data were first visualised as an intensity-normalised heatmap, revealing distinct patterns for each stimulus (Figure 1B). The samples broadly clustered by culture conditions; negative controls (i.e. RPMI medium alone, or RPMI combined with the transfection reagent Lyovec), bacterial ligands (lipopolysaccharide [LPS], PeptidoGlycaN [PGN]), and most viral mimetics (polyinosinic:polycytidylic acid [Poly(I:C)], Lyovec/3’3’-cyclic GMP-AMP [cGAMP], Lyovec/5’-ppp double stranded [ds]RNA) showing distinct clusters of responses along the rows of Figure 1B.

In negative control culture supernatants, detectable cytokines were generally similar for RPMI and Lyovec/RPMI. These included platelet-derived growth factor (PDGF)-BB, monocyte chemoattractant protein (MCP)1, interleukin (IL)-8, vascular endothelial growth factor (VEGF)A, and interleukin-1 receptor antagonist (IL-1Ra). Compared to RPMI alone, Lyovec/RPMI induced interferon (IFN)α and C-X-C Motif Chemokine Ligand 10 (CXCL10) in some children, and IL-1Ra and VEGFA were slightly increased on average (Supplementary Figure 1).

Cytokine production following ligand stimulation generally showed marked inter-individual variation, with some responses varying across orders of magnitude (Supplementary Figure 1, prominently e.g., IL-6 in response to LPS). Furthermore, the strength of the responses differed according to the stimulus. For example, as seen in Figure 1B, Tumour necrosis factor (TNF^23^), IL-10, IL-6, and IL-1β were strongly induced by LPS, PGN, and R848 (resiquimod), whereas IFNα, IFNγ, and CXCL10 were predominantly produced in response to Lyovec/cGAMP and Poly(I:C) in addition to R848. VEGF-A and IL-8, and to a lesser extent also PDGF-BB were – in contrast to interferons and CXCL10 – most strongly produced in response to PGN, LPS, and dsRNA.

Principle component analysis confirmed the clear separation of the responses dependent on the specific stimulus. In principal component analyses (PCA), Lyovec/RPMI and RPMI alone were largely superimposed indicating similar overall responses (Figure 1C-D). R848 was separated the furthest from RPMI across both PC1 and PC2, reflecting the potency of this stimulus, but the separation along PCs differed for the other stimuli. LPS, PGN, and Lyovec/dsRNA showed the strongest shift from control samples along PC1, whereas Poly(I:C) Lyovec/cGAMP separated from RPMI on PC2. For PC3, Lyovec/dsRNA and Poly(I:C) diverged from the other stimuli.

In summary, we observed considerable inter-individual variation in whole blood cytokine responses to innate stimuli. Each stimulus provoked a distinct response, aligning with expectations based on previous literature from adults^6^.

### Common genetic variants explain substantial variation in cytokine responses in early childhood

We first considered genetic determinants of variation in innate immune cytokine responses (Figure 2A). We performed cytokine quantitative trait loci (cyQTL)-mapping to investigate the association between genetic variants (single nucleotide polymorphisms, SNPs) and cytokine responses for cytokine-stimulus combinations that were within quantifiable range in at least 50% of the samples (see also Supplementary Figure 1). These analyses were performed using data from a subset of children due to missingness of covariate data (n = 259 [90.6% of total sample]; see methods for details). We set three thresholds for statistical significance: (i) a lenient screening threshold (*p* < 5x10^-6^); (ii) the commonly used ‘genome-wide’ significance (*p* < 5x10^-8^); and (iii) ‘study-wide’ significance for which the genome-wide significance threshold was made more strict to account for the number of effective comparisons^24,25^ (here: determined to be *p* < 1.39x10^-9^).

**Figure 2:**
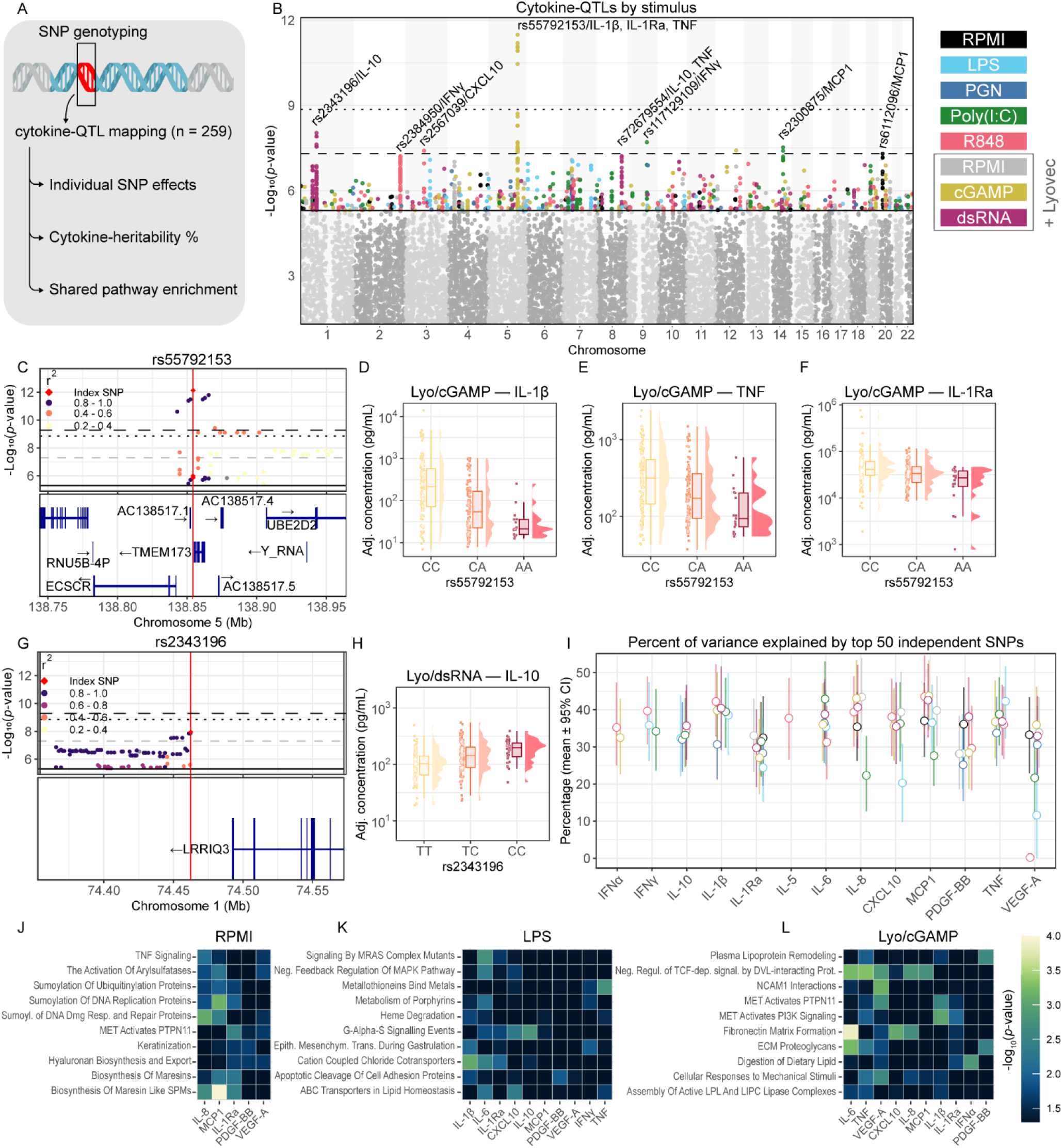
Genetic influences on innate immune cytokine responses. (A) overview of the analyses^132^. (B) Manhattan plot of cytokine-QTLs coloured by stimulus. The dashed lines indicate different levels of statistical significance: lenient screening threshold (bottom line; *p* < 5x10^-6^), genome-wide significance (middle line; *p* < 5x10^-8^), and study-wide significance (top line; *p* < 1.39x10^-9^). (C) Locuszoom plot (top) and genetrack (bottom) of rs55792153. (D-F) Covariate-adjusted concentrations of respectively IL-1β, TNF, and IL-1Ra upon stimulation with Lyovec/cGAMP, in children with different genotypes on this location. (G) Locuszoom (top) and genetrack (bottom) of rs2343196. (H) Covariate-adjusted concentrations of IL-10 upon stimulation with Lyo/dsRNA, in children with different genotypes on this location. (I) Graph indicating the percentage of variance explained by the top 50 LD-independent SNPs for each stimulus-cytokine combination. The graph is arranged by cytokine with colours indicating the stimuli (same colour legend as B). Dots indicate the estimated mean percent of variance explained, with whiskers indicating the 95% confidence interval. (J-L) Top 10 Reactome pathways that were enriched (nominal *p* < 0.05), shared between the highest proportion of cytokines for stimulation with each of respectively RPMI, LPS, and Lyovec/cGAMP. In this figure, covariates included in the models are time in freezer, exact incubation time, sex, seasonal infection burden, and granulocyte percentage; except panel I for which the underlying cytokine data was only adjusted for time in freezer and exact incubation time (as to not overestimate percent of *biological* variance explained by genetics). Boxplots are in the style of Tukey.

A single locus, rs55792153, met the study-wide significance threshold (Figure 2B). This SNP is located immediately downstream of *TMEM173*, which encodes stimulator of interferon genes (STING), the receptor for cGAMP (Figure 2C). Compared to major allele homozygotes (CC), those with an AA or heterozygous genotype produced lower IL-1β, IL-1Ra and TNF following stimulation with Lyovec/cGAMP (Figure 2D-F). Most other SNPs meeting or approaching the genome-wide (but not study-wide) significance threshold were not located near canonical immune-related genes. The second-most significantly associated SNP was rs2343196, located downstream of *LRRIQ3*, which is mainly expressed in the testes but also in many immune cells^26,27^ (Figure 2B, 2G). Compared to reference allele (TT) homozygotes, children heterozygous or homozygous for the variant allele C produced less IL-10 on stimulation with Lyovec/dsRNA (Figure 2H). Additional SNPs with or approaching genome-wide significance are shown in the supplementary data and include rs72679554 (near RP11-536K17.1, Supplementary Figure 2A-C), rs2384950 (strong linkage disequilibrium [LD] with a dense gene cluster; Supplementary Figure 2D-E), rs2300875 (in *ACTN1*, Supplementary Figure 2F-G), and rs6112096 (nearest to *DTD1*; Supplementary Figure 2H-I). Together, these data reinforce that specific SNPs markedly impact specific stimulus-cytokine combinations (or sometimes more broadly the cytokine response, depending on the locus).

We next investigated the total variance in cytokine production that could be explained by SNPs. To avoid overfitting given the large number of stimulus-output combinations relative to the sample size, we focused on the top 50 (LD-independent, by *p*-value) associated SNPs per stimulus-cytokine combination. Overall, genetic variation appears to be a strong determinant of cytokine responses in preschool children; the combination of the top 50 SNPs explained approximately 20-45% of inter-individual variation for most cytokine-stimulus combinations (Figure 2I). MCP1 responses to Lyovec/cGAMP and R848 stimulation showed the highest percent of variance explained (respectively 43.7% [95%CI: 34.6 – 53.3] and 43.5% [95%CI: 33.3 – 54.6], Figure 2I). In contrast, for LPS- or R848-stimulated levels of VEGF-A, almost none of the variation was explained by the top 50 SNPs.

We then performed gene-set (Reactome) enrichment analyses of SNPs within a window of 35 kb upstream to 10 kb downstream of genes to investigate potential shared pathways across stimulus-cytokine combinations. Pathways enriched for SNPs associated with baseline cytokine production included “Biosynthesis of maresins”/”Biosynthesis of maresin-like Specialised Pro-resolving Mediators” (RPMI) and FLT3-related gene-sets such as “FLT3 signaling through Src family kinases” (Figure 2J, Supplementary Figure 2J).

Across the different stimuli, some well-understood signalling pathways were frequently enriched. Chloride transporter-related pathways were shared between LPS and R848 (Figure 2K, Supplementary Figure 2J). PI3K-related signalling (e.g., “Activated NTRK2 signals through PI3K”, “MET activates PI3K signaling”) was shared between PGN and Lyovec/cGAMP (Figure 2L, Supplementary Figure 2J). The cytokines IL-1β, IL-6, TNF, IL-10, and IL-1Ra often shared pathways within a stimulus, highlighting their central roles in innate immune responses and suggesting that their expression is co-regulated to a degree.

### Dimensionality reduction using CytoMod – Cytokine co-expression modules identify recurring response patterns

We next investigated additional host factors and environmental determinants of childhood cytokine responses. First, we used a freely available python module, Cytomod^28^, to reduce the number of dimensions and hence the multiple testing burden. Briefly, CytoMod clusters cytokines into ‘modules’ based on patterns of co-expression (Figure 3A). The modules are determined by unsupervised clustering of cytokines after adjusting for the participant-level mean cytokine value. The final module composition is then based on pairwise reliability scores using a bootstrapping approach. Module expression scores are calculated by taking the mean of standardised cytokine concentrations included in that module^28^.

**Figure 3:**
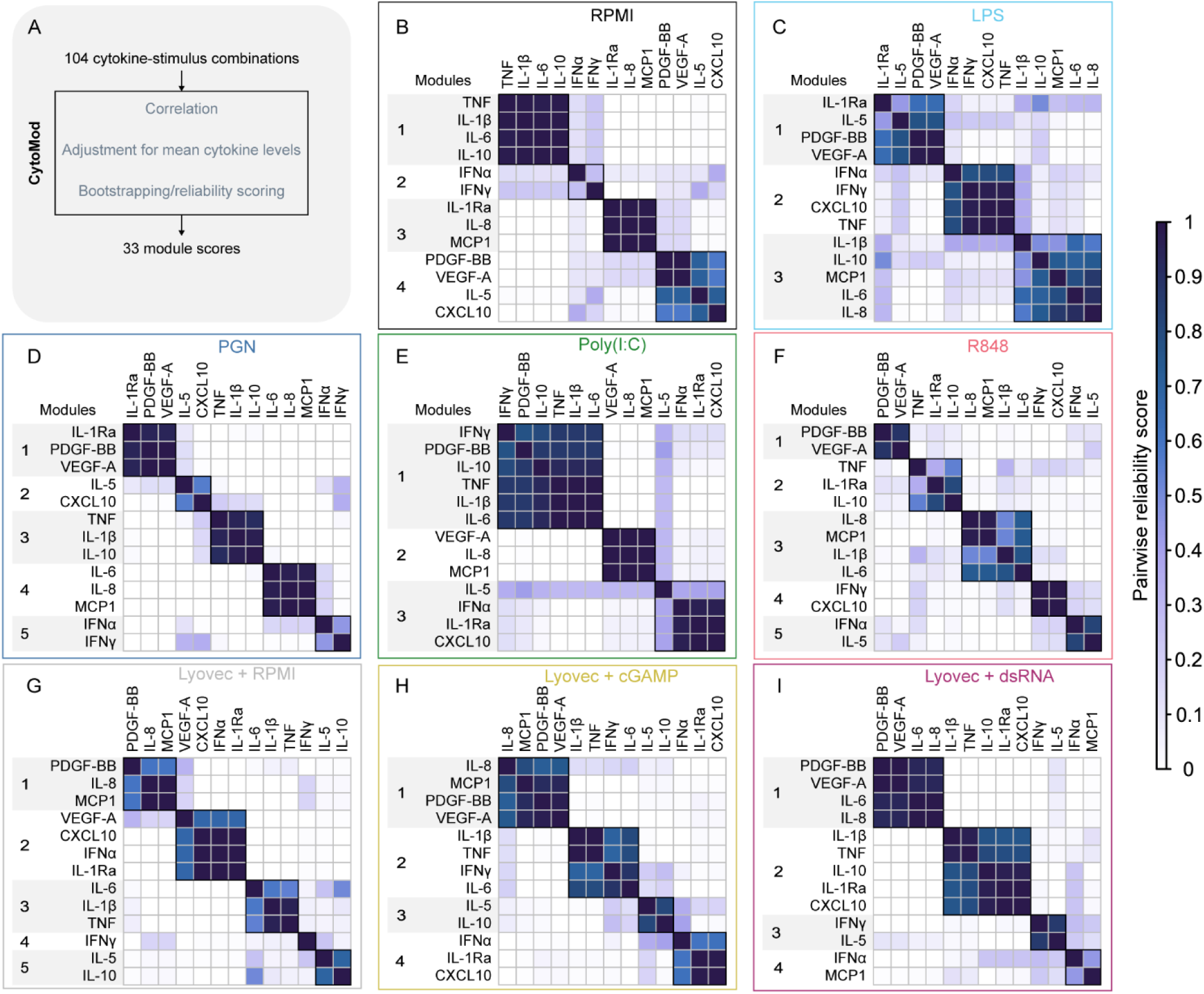
Establishing modules of co-expressed cytokines. (A) Schematic overview of the CytoMod procedure. Clustered heatmaps of CytoMod pairwise reliability score, annotated with the cytokine modules for the stimuli: (B) RPMI, (C) LPS, (D) PGN, (E) Poly(I:C), (F) R848, (G) Lyovec/RPMI, (H) Lyovec/cGAMP, (I) Lyovec/dsRNA.

We performed the clustering for each stimulus separately (Figure 3B-I), as each stimulus affects expression of cytokines differentially. In these analyses, CytoMod assigned between 3 and 5 cytokine modules per stimulus (Table 2), reducing the number of parallel comparisons from 104 stimulus-cytokine combinations to 33 cytokine module expression scores. The correlation between cytokines within each module, as well as between each cytokine and its module expression score are shown in Supplementary Figure 3. While the cluster composition differed between stimuli, some general patterns were evident; TNF, IL-6, IL-1β, and IL-10 responses tended to cluster together, as did IFNα and IFNγ, as well as IL-8, MCP-1, VEGFA, and PDGF-BB. For each stimulus, cytokines that were or were not released also tended to cluster together.

**Table 2:**
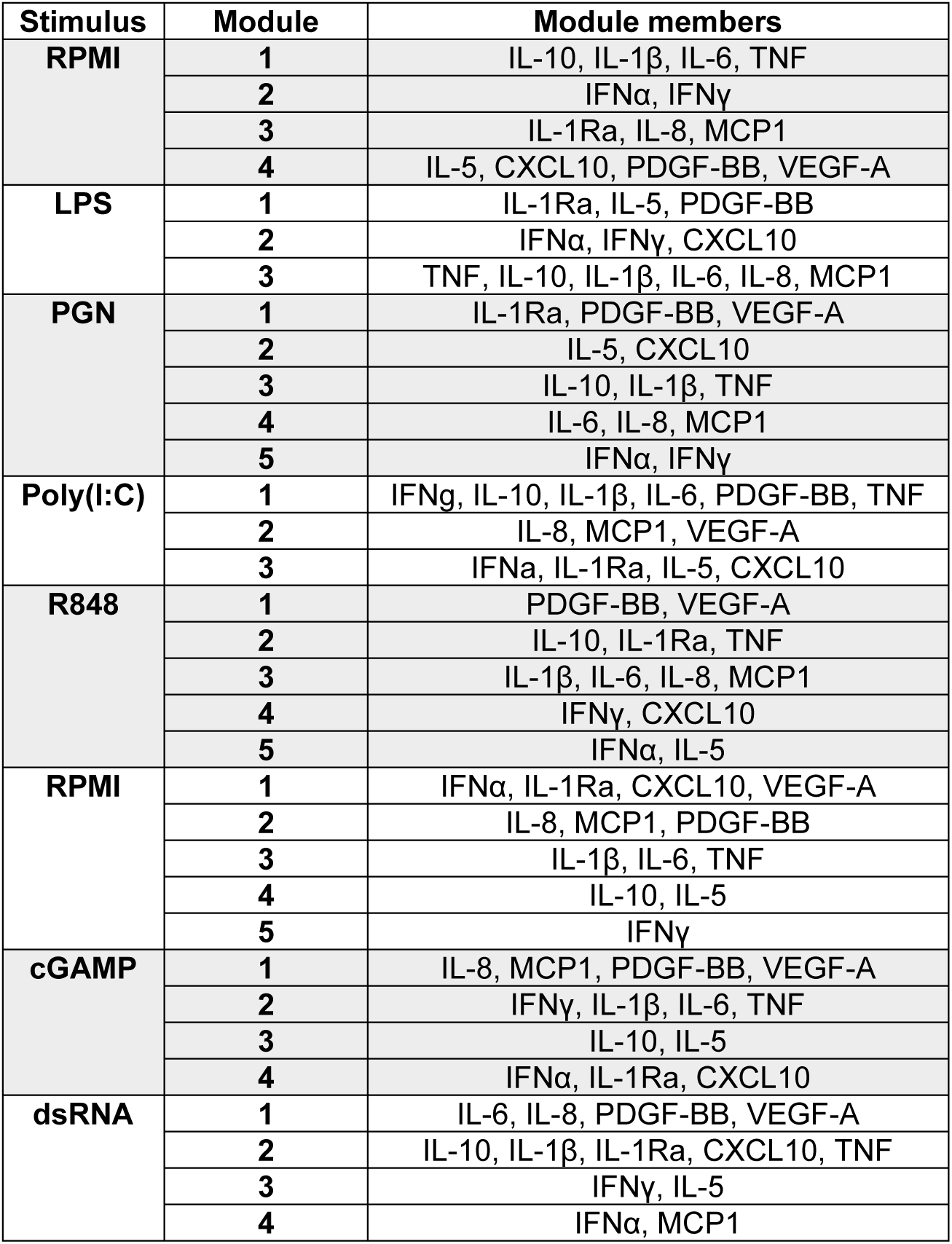
Members of each cytokine module.

| Stimulus | Module | Module members |
| --- | --- | --- |
| RPMI | 1 | IL-10, IL-1β, IL-6, TNF |
|  | 2 | IFNα, IFNγ |
|  | 3 | IL-1Ra, IL-8, MCP1 |
|  | 4 | IL-5, CXCL10, PDGF-BB, VEGF-A |
| LPS | 1 | IL-1Ra, IL-5, PDGF-BB |
|  | 2 | IFNα, IFNγ, CXCL10 |
|  | 3 | TNF, IL-10, IL-1β, IL-6, IL-8, MCP1 |
| PGN | 1 | IL-1Ra, PDGF-BB, VEGF-A |
|  | 2 | IL-5, CXCL10 |
|  | 3 | IL-10, IL-1β, TNF |
|  | 4 | IL-6, IL-8, MCP1 |
|  | 5 | IFNα, IFNγ |
| Poly(I:C) | 1 | IFNγ, IL-10, IL-1β, IL-6, PDGF-BB, TNF |
|  | 2 | IL-8, MCP1, VEGF-A |
|  | 3 | IFNα, IL-1Ra, IL-5, CXCL10 |
| R848 | 1 | PDGF-BB, VEGF-A |
|  | 2 | IL-10, IL-1Ra, TNF |
|  | 3 | IL-1β, IL-6, IL-8, MCP1 |
|  | 4 | IFNγ, CXCL10 |
|  | 5 | IFNα, IL-5 |
| RPMI | 1 | IFNα, IL-1Ra, CXCL10, VEGF-A |
|  | 2 | IL-8, MCP1, PDGF-BB |
|  | 3 | IL-1β, IL-6, TNF |
|  | 4 | IL-10, IL-5 |
|  | 5 | IFNγ |
| cGAMP | 1 | IL-8, MCP1, PDGF-BB, VEGF-A |
|  | 2 | IFNγ, IL-1β, IL-6, TNF |
|  | 3 | IL-10, IL-5 |
|  | 4 | IFNα, IL-1Ra, CXCL10 |
| dsRNA | 1 | IL-6, IL-8, PDGF-BB, VEGF-A |
|  | 2 | IL-10, IL-1β, IL-1Ra, CXCL10, TNF |
|  | 3 | IFNγ, IL-5 |
|  | 4 | IFNα, MCP1 |

Throughout the following sections, we present associations between variables of interest and the expression scores of cytokine modules as primary analysis, with *p*-values adjusted for multiple testing using the FDR method. As this is a data-scarce field, in secondary analyses we report associations between variables of interest and individual stimulus-cytokine combinations with nominal *p* < 0.05.

### Limited associations between child characteristics and cytokine responses at four years of age

We examined associations between exact age, sex, and measures of adiposity (measured at blood collection) and cytokine module expression (Figure 4A). For these analyses we used data from 286 children (136 girls and 150 boys; Figure 4B) approximately four years of age (range: 3.9 – 5.6 years; median: 4.2 years; Figure 4C). Anthropometric and adiposity measures (BMI [n = 286] and body fat percentage [n = 269]) were mostly within the normal range for their sex and age^29^ (Figure 4D, E). After FDR correction, we found no evidence of associations between these anthropometric measurements and cytokine module expression (Figure 4F). Given the data scarcity in this field, we additionally reported nominal *p*-values < 0.05 for these analyses (Figure 4F-I).

**Figure 4:**
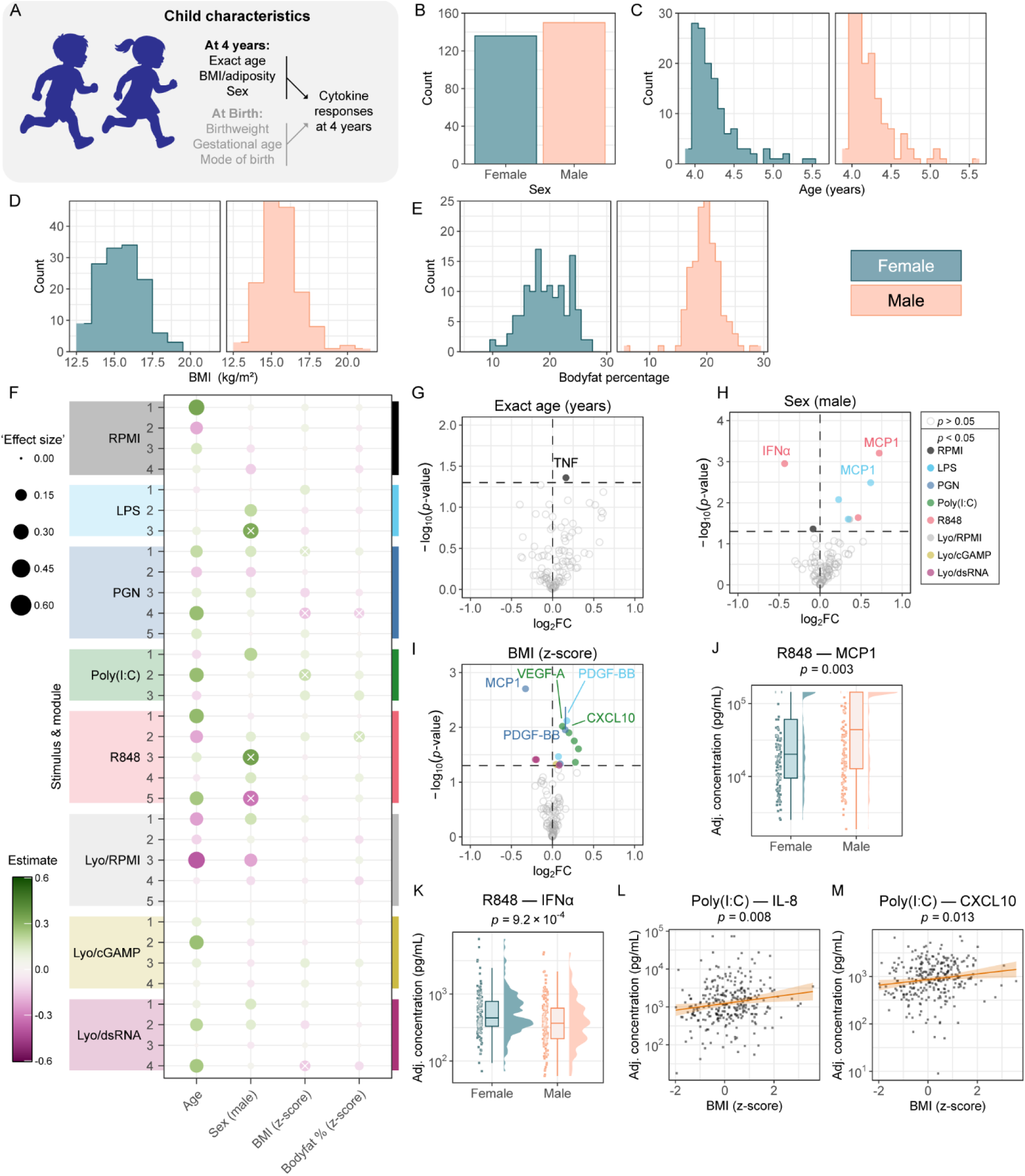
Anthropometrics and cytokine responses at four years of age. (A) Schematic description of analyses in this figure. (B) Number of girls and boys in these analyses. (C) Distribution of age. (D-E) Distribution of measures of adiposity: BMI and bodyfat percentage. (F) Dotplot summarising regression results between each variable of interest and cytokine module expression score. In each dot ‘x’ indicates nominal *p* < 0.05. ‘Effect size’ was calculated as absolute value of the regression estimate. (G) Volcano plot indicating change per 1 year increase in age. (H) Volcano plot indicating the difference between boys and girls (right: higher in boys, left: higher in girls). (I) Volcano plot indicating change per 1 SD increase in BMI. (J) Covariate-adjusted concentrations of MCP1 following R848 stimulation, in male children versus female children. (K) Covariate-adjusted concentrations of IFNα following R848 stimulation, in boys versus girls. (L) Covariate-adjusted concentrations of IL-8 following poly(I:C) stimulation, as a function of BMI z-score. (M) Covariate-adjusted concentrations of CXCL10 following poly(I:C) stimulation, as a function of BMI z-score. In this figure, ‘covariate adjusted’ includes time in freezer, exact incubation time, and sex, age, and BMI (except when one of these was the variable of interest). The *p*-values in panels G-M are unadjusted. The ribbon surrounding the regression line in panels L-M indicates 95% confidence interval of the slope. Boxplots are in the style of Tukey.

We did not find evidence that exact age (within the narrow available age-range, Figure 4C) was associated with any of the cytokine responses (Figure 4F, G). Sex differences in immune responses are well-described in adults (see for example references ^3,6,30–33^), but we found only a small number of differences in this cohort: modules LPS-3 and R848-3 were higher in male children, whereas R848-5 was higher in female children (Figure 4F). These differences were driven by MCP1 (included in both LPS-3 and R848-3; Figure 4H, J) and IFNα (included in R848-5; Figure 4H, K), respectively – well-known examples from adult studies^32,33^. We found evidence that measures of adiposity were associated with cytokine responses, although the effect size was small in this cohort of predominantly normal-weight children. The BMI z-score (derived from WHO standards data^29^) was positively associated with PGN-1 levels and inversely with PGN-4. There was also a negative association between BMI z-score and Lyovec/dsRNA-4. Finally, the module Poly(I:C)-2 as well as more broadly individual cytokine responses to poly(I:C) were associated with BMI z-score (Figure 4F, I, L, M). Body fat percentage showed weak evidence of associations patterns, similar to BMI z-score (Figure 4F).

### Pregnancy and perinatal variables show weak or non-significant associations with cytokine responses

As pregnancy and perinatal factors may influence immune development^16,34^, we investigated associations between mode of birth, birth weight, and gestational age and cytokine modules (Figure 5A). Most children were born at term (37 to 42 weeks gestational age; median: 39.6 weeks, range: 32.1 – 41.9 weeks; Figure 5B), and with a birth weight within the normal range (median: 3.53 kg, range: 1.61 – 5.41; Figure 5C). Of the 286 children included in these analyses, 102 (35.6%) were born by caesarean section (Figure 5D).

**Figure 5:**
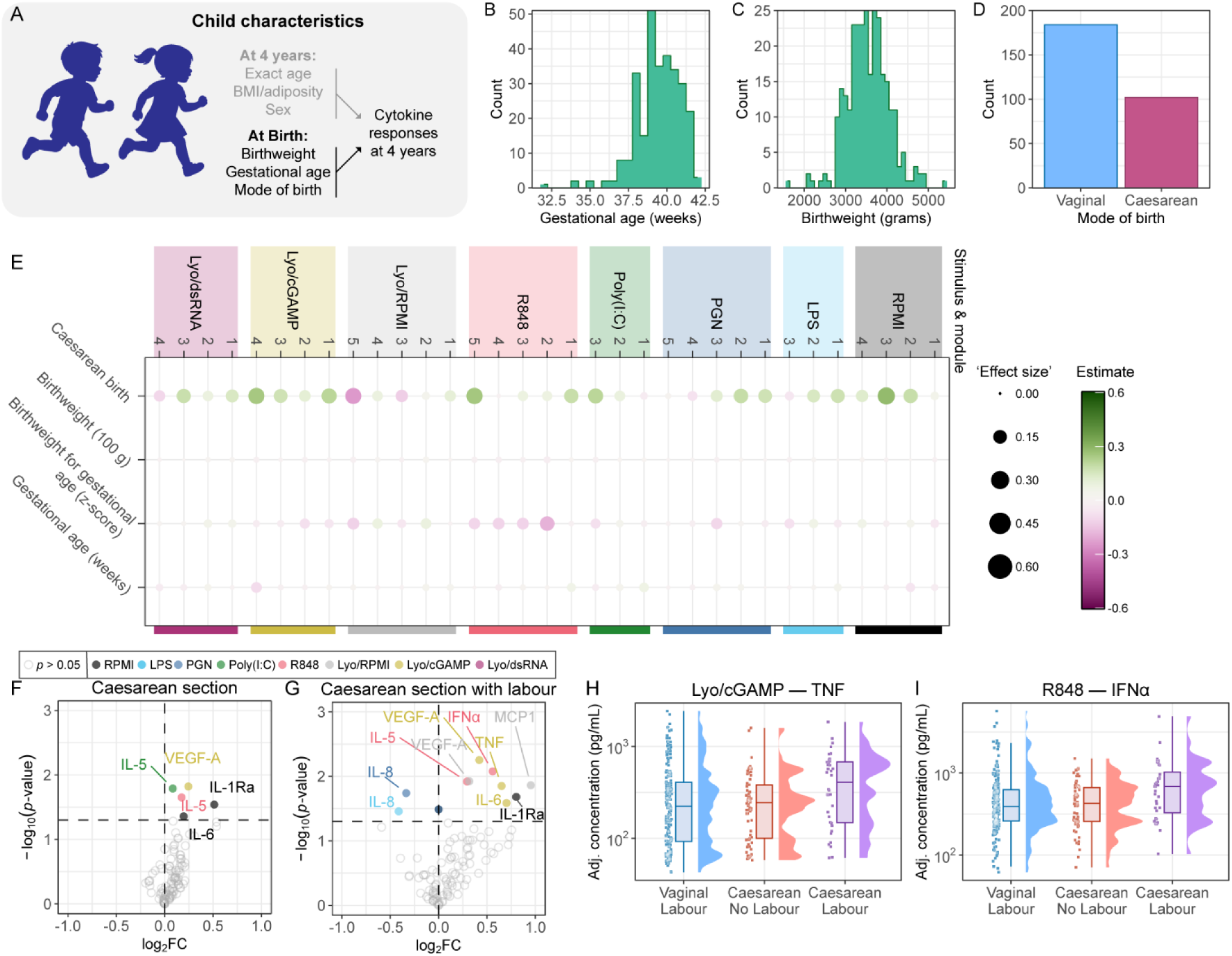
Pregnancy and perinatal variables and cytokine responses at four years of age. (A) Schematic description of analyses in this figure. (B) Distribution of gestational age. (C) Distribution of birthweight. (D) Proportion of children born via vaginal birth and caesarean section. (E) Dotplot summarising regression results between each variable of interest and cytokine module expression score. ‘Effect size’ was calculated as absolute value of the regression estimate. (F) Volcano plot indicating the difference between birth modes (right: higher in caesarean section, left: higher in vaginal birth). (G) Volcano plot indicating the difference according to having experienced labour prior to caesarean section (reference: children who had not experienced labour before caesarean section). (H) Covariate-adjusted concentrations of TNF following Lyovec/cGAMP stimulation, stratified by differences in mode of birth. (I) Covariate-adjusted concentrations of IFNα following R848 stimulation, stratified by differences in mode of birth. In this figure, ‘covariate adjusted’ includes time in freezer, exact incubation time, and sex, age, and BMI. The *p*-values in panels F-G are unadjusted. Boxplots are in the style of Tukey.

We found no statistically significant associations between cytokine module expression and birthweight or gestational age when analysed separately, nor for the birthweight z-score derived per sex and accounting for gestational age^35^ (Figure 5E). We found weak evidence for associations between these exposures and production of individual cytokines (Supplementary Figure 4A-C).

We also found no statistically significant associations between mode of birth and cytokine module expression. However, we did observe that the model estimates were mostly positive, suggesting that cytokine modules were generally higher in children born by caesarean section (Figure 5E). This is also visualised at the level of individual cytokines in Figure 5F, as a right-skewed volcano plot. We hypothesised that there may be a subgroup of children whose cytokine responses were more strongly associated with caesarean birth than others. We therefore investigated if the experience of labour, an inflammatory process^36^, prior to caesarean section was associated with differences in cytokine module expression or individual cytokine responses. We observed that, while the effects were statistically non-significant, there was some evidence that the pattern of differences in cytokine modules as well as individual cytokines were more pronounced in children born via caesarean section after experiencing labour (n = 36; Figure 5G [caesarean section with labour versus vaginal birth], Supplementary Figure 4D [subgroup comparisons]). The evidence was strongest for modules R848-5 and Lyo/cGAMP-1, which include IFNα (Figure 5H) and TNF (Figure 5I), respectively. Given the limited sample size of caesarean subgroups, these analyses should be interpreted as exploratory.

### Systemic inflammation and leukocyte composition are strongly associated with cytokine production capacity

We next investigated how inflammatory biomarkers, frequently assayed in cohort studies in favour of more labour-intensive stimulation assays, are related to whole blood cytokine production (Figure 6A). We considered three plasma measures of systemic inflammation; high-sensitivity C-reactive protein (hsCRP, n = 280), a commonly measured biomarker of acute systemic inflammation^37^; glycoprotein acetyls (GlycA, n = 283), a composite biomarker that is more stable than hsCRP and better captures chronic systemic inflammation^38–40^; and granulocyte-to-lymphocyte ratio (GLR, n = 265), which is increased during acute inflammation and trained immunity due to increased granulocyte production^41,42^. Forty-seven (16.8%) of the children had hsCRP level concentrations that were below the limit of detection (0.001 µg/mL), whereas all GlycA concentrations were within the quantifiable range (Figure 6B). GLR varied substantially across the population (median 1.44, range: 0.44 – 4.46; Fig 6C-D, Supplementary Figure 5A-C). All three biomarkers were log-transformed and internally standardised for statistical analyses.

**Figure 6:**
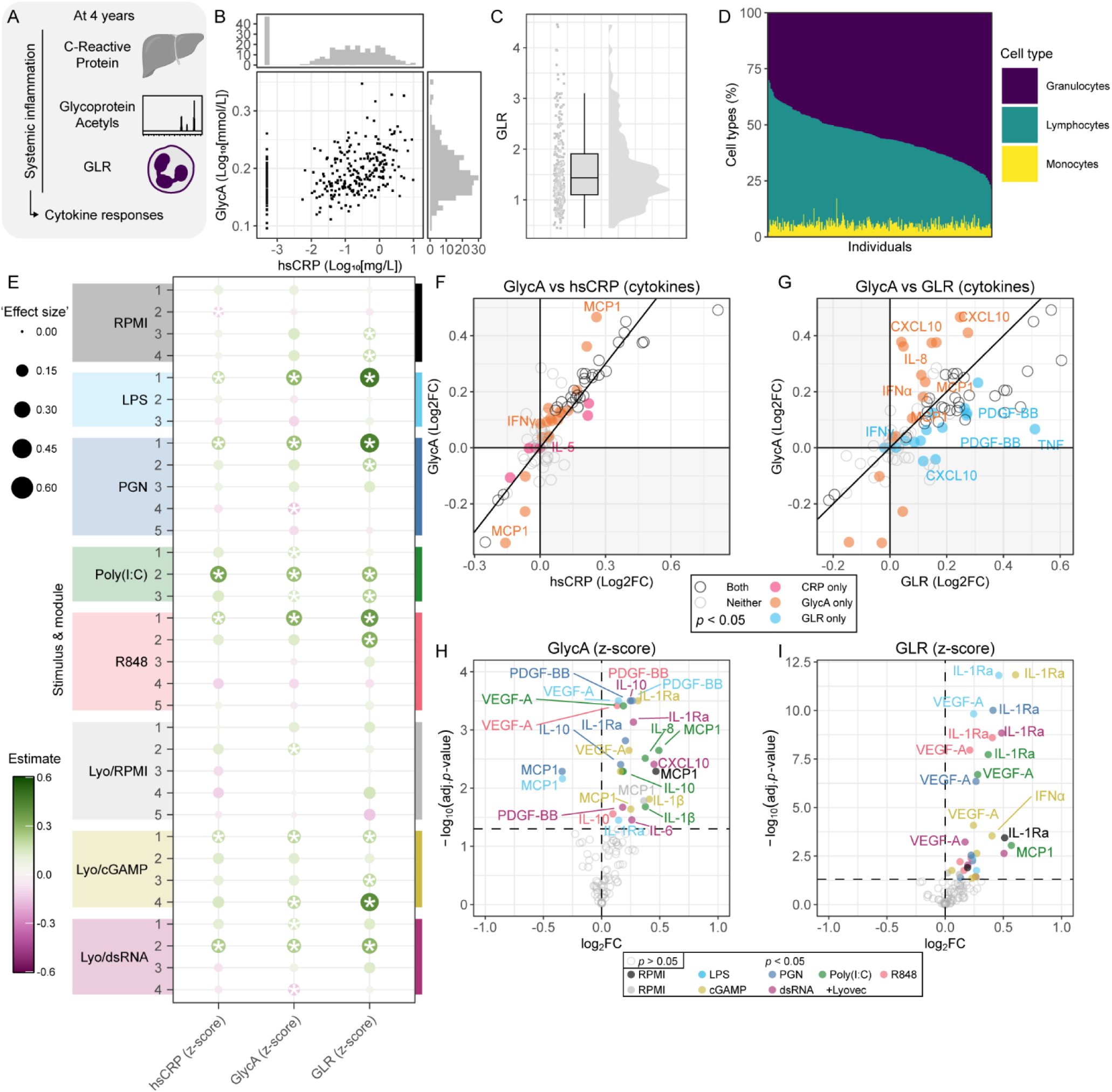
Associations between markers of systemic inflammation and cytokine responses. (A) Schematic overview of the performed analyses. (B) Distribution of GlycA and hsCRP concentrations plotted against each other. (C) Distribution of the granulocyte-to-lymphocyte ratio. (D) Relative proportion of major leukocyte types, ordered by granulocyte percentage. (E) Dotplot summarising regression results between each variable of interest and cytokine module expression score. In each dot ‘*’ indicates FDR-adjusted *p* < 0.05. ‘Effect size’ was calculated as absolute value of the regression estimate. (F) Comparative plot indicating the average fold change in cytokine levels when GlycA or hsCRP is increased by 1 SD. The points are coloured by the indicated nominal *p*-value significance categories. (G) Comparative plot indicating the difference change in cytokine levels when GlycA or GLR is increased by 1 SD. The points are coloured by the indicated nominal *p*-value significance categories. For panels F-G, points in the white areas are concordant between measures; points in the grey areas are discordant. (H) Volcano plot indicating the change in cytokine levels with an increase of 1 SD in GlycA. (I) Volcano plot indicating the change in cytokine levels with an increase of 1 in GLR. In this figure, ‘covariate adjusted’ includes time in freezer, exact incubation time, and sex, age, and BMI. The *p-*values in panels F-G are unadjusted. Boxplots are in the style of Tukey. GLR, granulocyte-to-lymphocyte ratio.

In total, associations between any of the three inflammatory markers and the various cytokine modules were overwhelmingly positive: we found 3 negative associations and 30 positive associations (Figure 6E). We observed considerable overlap between the 7 cytokine modules associated with hsCRP and the 13 modules associated with GlycA (Figure 6E). GlycA was also associated with more individual cytokines (Figure 6F). Only a single module (RPMI-2) was (negatively) associated with hsCRP but not GlycA (Figure 6E, Supplementary Figure 5D). Generally, an increase of 1 standard deviation in GlycA levels was associated with a larger increase or decrease in module expression score, compared to a similar increase in hsCRP (Supplementary Figure 5D).

GLR showed the strongest associations overall and was associated with 13 cytokine modules, more than GlycA (Figure 6E, 6G, Supplementary Figure 5E). Both biomarkers were often associated with modules containing PDGF-BB, VEGF-A, and IL-1Ra. However, while most of the modules overlapped between GlycA and GLR, different cytokines within these modules drove these associations. We therefore observed that a substantial number of stimulus-cytokine pairs were either associated with GLR only, or only with GlycA. For stimulus-cytokine pairs that were associated with both biomarkers, associations were more evident for GLR (Figure 6G). The associations with GLR were strongly driven by IL-1Ra and VEGF-A, whereas GlycA was linked with PDGF-BB and a mixture of other cytokines (Figure 6H-I).

In summary, we found that all three markers of inflammation were associated with cytokine module expression, but to different extents and that associations were driven by different individual cytokines.

As GLR showed the strongest association with any given module, we explored associations between cytokine responses and relative abundance of lymphocytes, monocytes, and granulocytes (predominantly neutrophils). Compared to GLR, percentage of granulocytes showed greater evidence of associations with cytokine modules that incorporate VEGF-A and IL-1Ra (Supplementary Figure 5F, G). As anticipated given the inverse relationship between granulocyte and lymphocyte percentages, this was mirrored by the lymphocyte percentage (Figure 6B, Supplementary Figure 5F, H). The percentage of monocytes was more stable across participants, when considering percentage-point differences (Figure 6D, Supplementary Figure 5C), and was associated (at the individual cytokine level) with production of classical monocyte cytokines such as IL-6 and IL-1β, and also IFNα (Supplementary Figure 4I). For additional context, it is important to mention that the IL-1β/IL-1Ra ratio (which has been proposed as a measure of biological IL-1β activity^43,44^) was low across stimuli (Supplementary Figure 1).

### Seasonal community viral infection burden coincides with heightened antiviral cytokine responses

We next investigated the seasonality of cytokine responses in this cohort (Figure 7A)^6,8,45^. Cytokine modules in response to particularly Lyovec/cGAMP and Lyovec/dsRNA peaked in the Southern Hemisphere winter (June-August, Supplementary Figure 6). Other stimuli showed a similar pattern, although not for all cytokine modules. The individual cytokines IL-1Ra and CXCL10 showed the most consistent seasonal pattern across stimuli (Supplementary Figure 7). Of note, the sampling density over time decreased towards the end of the study period, as reflected by the widening of the confidence bands (Supplementary Figures 6 and 7).

**Figure 7:**
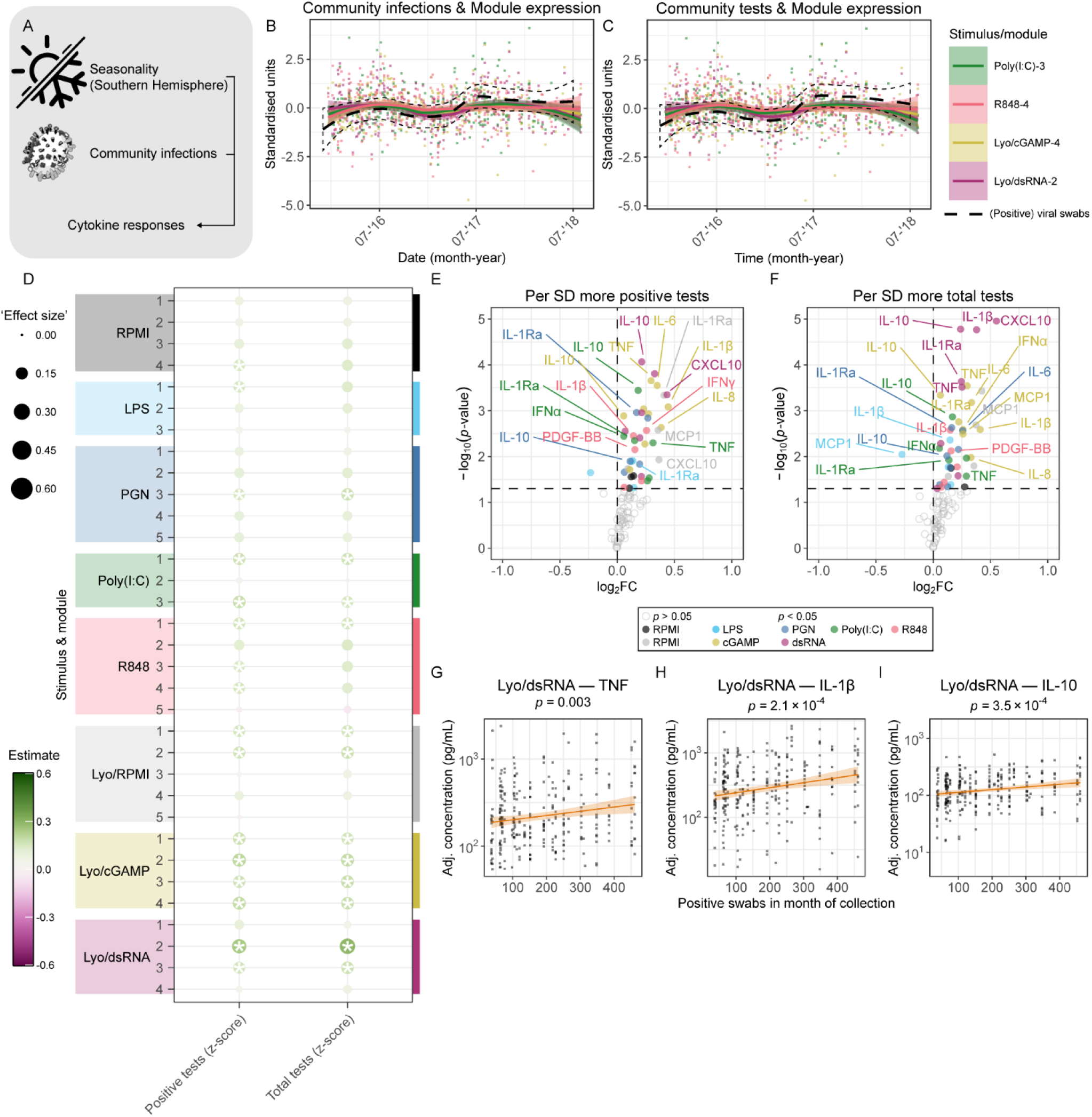
Seasonal community infections coincide with peaks in cytokine responses. (A) Schematic overview of the performed analyses^133^. (B) Number of positive viral swabs (log-transformed and standardised) overlaid on expression scores of modules that exhibit seasonal variation. (C) Same as for B but showing the total number of swabs taken. (D) Dotplot summarising regression results between each variable of interest and cytokine module expression score. In each dot ‘*’ indicates FDR-adjusted *p* < 0.05. ‘Effect size’ was calculated as absolute value of the regression estimate. (E) Volcano plot indicating the change in cytokine levels with an increase of 1 SD in positive viral swabs in the month of blood collection. (F) Volcano plot indicating the change in cytokine levels with an increase of 1 SD in total viral swabs in the month of blood collection. (G-I) Scatter plots showing covariate-adjusted concentrations of respectively TNF, IL-1β, or IL-10 upon stimulation with Lyovec/dsRNA, as a function of positive viral swabs in the month of blood collection. In this figure, ‘covariate adjusted’ includes time in freezer, exact incubation time, and sex, age, and BMI. The *p-*values in panels E-I are unadjusted. The ribbon around the regression line (panels B-C, G-I) indicates the 95% confidence interval around the estimate.

We hypothesised that the higher incidence of viral infections in winter – a common cause of morbidity in young children^46–48^ – may contribute to the seasonal variation in cytokine responses. We leveraged ecological infection data from Snotwatch^49^ (NB: independent of BIS) to investigate the association between circulating viral infections and cytokine responses. We quantified the number of positive respiratory viral swabs (Supplementary Figure 8A), the total number of viral swabs collected (a proxy for how many people experienced symptoms of viral infections that warranted a swab; Supplementary Figure 8B), and the positivity rate (Supplementary Figure 8C) for each month in the BIS sample collection date interval (spanning December 2015 through July 2018). Taking four cytokine modules that show seasonal differences as an example, we overlaid the expression of modules Poly(I:C)-3, R848-4, Lyovec/cGAMP-4, and Lyovec/dsRNA-2 with the standardised, log-normalised number of positive viral swabs in a given month. We observed a similar, though imperfect, pattern of peaks in childhood infection and expression of these modules (Figure 7B). We observed a similar pattern for the total number of swabs taken (Figure 7C).

We then studied the associations between (ecological, population-level) community infections in the month of blood collection and cytokine production. We found that 14 cytokine modules were associated with the number of positive viral swabs in the month of blood collection (Figure 7D). The associations with total number of tests in the month of blood collection was somewhat weaker, both in terms of regression estimates and number of significantly associated modules (Figure 7D). Congruent with the modules showing the strongest seasonality, the pattern of association was strongest for Lyovec/cGAMP and Lyovec/dsRNA. We found that individual cytokines particularly in response to viral stimuli were associated with the number of (positive) tests. This included cytokines such as IFNα and CXCL10, which are typically associated with antiviral responses, but also prominently IL-1β, IL-1Ra, IL-10, and TNF. This suggests a broadly increased inflammatory response, rather than a targeted antiviral mechanism.

## Discussion

In this population-derived cohort of children around 4 years of age, we demonstrated substantial inter-individual variation in early life cytokine responses and quantified the relative contributions of genetic, host-intrinsic, and selected environmental determinants. Genetic variants explained the largest proportion of variation in cytokine production capacity, followed by strong associations with seasonal viral infections and cell type composition. Biomarkers of inflammation associated with cytokine responses in marker-specific patterns, likely reflecting their underlying origins. We found some evidence that mode of birth in combination with the experience of labour may have long term effects on cytokine responses.

We identified several genomic loci associated with cytokine production capacity. The strongest association mapped to the *TMEM173*/STING locus, where variants were linked to a marked decrease in production of IL-1β, TNF, and IL-1Ra. Direct comparison with adult cyQTL studies was not possible as these have not used cGAS/STING-specific stimuli. However, follow-up studies are warranted given the central role of the cGAS-STING pathway in host defence, and evasion of STING being a common pathogenic mechanism^50^.

Unlike cohorts of Dutch descent^7,24,25^, we did not identify cyQTLs in the *TLR1-TLR6-TLR10* cluster. This could reflect ancestry differences as the *TLR1-TLR6-TLR10* locus differs markedly between populations due to introgression events^51–53^. A paediatric cohort from Tanzania, where these introgressions did not take place, also did not identify this as a key locus^21^. An age-related effect is unlikely but cannot be excluded as both cohorts that did not identify this locus are paediatric. Overall, the top 50 LD-independent SNPs explained a substantial proportion (∼20-45%) of variance in cytokine production. Compared to the most analogous study in adults^8^, we found comparable or higher percentages of variance explained – but in a whole blood stimulation assay, which inherently has more non-genetic variation than the PBMC-stimulations^54^ reported in adults. This aligns with the genetic influence on cytokine responses decreasing with age^13^, which likely reflects the increasing cumulative exposure to each individual’s uniquely diverging environments. A more rigorous comparison with adult studies was not possible due to differences in experimental design (cell type, stimuli, analytes) and analytical approaches.

In contrast to adult data, sex-related differences in cytokine responses were modest, consistent with the pre-pubertal age of this cohort ^3,30,31,33,55–58^. This is unlikely to be explained by limited statistical power as several adult studies reporting sex differences were of similar or smaller size. We observed differences in MCP1 production (higher in male children), suggesting a genetic mechanism beyond the hormonal influences on circulating MCP1 levels in adults^33^. The observed increase in TLR7/8-driven IFNα responses in female children aligns with known X-linked regulation of antiviral sensing and supports that sex differences in innate immunity are detectable even in early life, albeit at smaller magnitude than in adulthood^56,59^.

There was some evidence that children delivered by caesarean section had stronger cytokine responses, particularly the subgroup that had been exposed to labour (presumably emergency caesarean sections). This aligns with evidence linking caesarean delivery with higher rates of childhood infection-related hospitalisation^60^ and increased risk of several inflammatory disorders^61^. Further studies incorporating clinical indications for caesarean delivery are needed to clarify these associations. As the associations at the module level were not significant after correction for multiple testing, and *p-*values for individual cytokines were unadjusted, these findings should be interpreted cautiously and require validated in independent cohorts.

Our findings regarding inflammation biomarkers reflect differences between cell-intrinsic cytokine production capacity and differences due to cellular composition. In related work, we have previously shown associations between GlycA and LPS/PGN-induced monocyte-associated cytokines^62^. Here, we show GlycA and GLR were both associated with various cytokine modules, but they did so through distinct cytokine signatures. Notably, IL-1Ra and VEGF-A were strongly associated with GLR and granulocyte abundance, the same cytokines that were relatively poorly explained by the top SNPs. This suggests that some cytokine responses are influenced more by cellular composition than (genetic) cell-intrinsic capacity, although both cell composition and ‘per-cell’ responsiveness are largely determined by the state of bone marrow progenitors^42,63^.

Indicators of respiratory viral infection incidence at a population level were associated with differences in cytokine responses in pre-school children, particularly responses to stimulation of antiviral pathways. For these analyses we considered positive PCR for any of the tested respiratory viruses, but it would be valuable to consider effects of specific viruses in appropriately sized follow-up cohorts. Nonetheless, cumulative or non-specific infection burden remains highly relevant: we have previously shown in BIS that total infection burden is associated with adverse plasma profiles of lipids and metabolites as early as 12 months of age^64^. These findings suggest that infection burden is a potentially modifiable determinant of long-term health^65^.

Together, these findings indicate that early childhood is a critical period during which innate immune responses are shaped strongly by genetic variation and to a more limited extent by the other host and exogenous environmental exposures considered in this study. A growing body of evidence suggests that childhood inflammation contributes to later cardiometabolic disease risk^66–69^, together with increasingly common^70,71^ traditional (and pro-inflammatory^72,73^) risk factors such as obesity and type 2 diabetes mellitus. Furthermore, infection burden during childhood is an emerging cardiovascular risk factor^64,74^. Understanding the determinants of variability in innate immune responses during this critical window is essential to identification of – and intervening in – children at risk before clinical disease emerges^69,75,76^.

## Limitations

We acknowledge a number of limitations. First, genetic analyses are traditionally performed in much larger cohorts to identify rarer variants or variants with smaller effect sizes. Nonetheless, prior studies^7,8,54,77^ suggest that moderate sample sizes can identify impactful genetic variants with larger effects. Second, the flow cytometry analyses used in this study were performed using very limited cell type markers, as they were designed at inception of this cohort when advanced methodology was not available at the study site. Third, the ecological, population-level estimate of respiratory viral infections was biased towards the Greater Melbourne area, rather than specifically in the Barwon area where the study was conducted. As the analysis of cytokines and viral exposure is ecological, the viral exposure data is not specific to participating children and their families. Further studies should also incorporate data on exposure to specific pathogens, and the range of pathogens should be extended to include other viral infections including gastrointestinal viruses. Finally, in this study, biomedical assessments were performed at approximately four years of age only, meaning we lacked a range of ages to identify meaningful effects of age difference, which could be an important determinant of immune responses in early childhood (especially considering that relatively modest age differences in young children constitute a larger proportion of lifetime than in adults). Similarly, BMI and body fat percentage were generally within a normal-range in this cohort, limiting our capacity to identify effects of these exposures on immune responses. We recognise that there is an immense number of potential environmental factors that may influence innate immune responses and that we have studied a small subset of those.

## Supporting information

Separate hires main figures

Supplementary complete model result tables

## Acknowledgements

We would like to thank all participants of the Barwon Infant Study, including their parents. Special thanks also go to the entire BIS team who enabled sample collection and provided essential laboratory support. The establishment work and infrastructure for the BIS was provided by the Murdoch Children’s Research Institute (MCRI), Deakin University and Barwon Health. Subsequent funding was secured from the National Health and Medical Research Council of Australia, The Jack Brockhoff Foundation, the Scobie Trust, the Shane O’Brien Memorial Asthma Foundation, the Our Women’s Our Children’s Fund-Raising Committee Barwon Health, The Shepherd Foundation, the Rotary Club of Geelong, the Ilhan Food Allergy Foundation, GMHBA Limited and the Percy Baxter Charitable Trust, Perpetual Trustees and the Minderoo Foundation. In-kind support was provided by the Cotton On Foundation and CreativeForce. Research at Murdoch Children’s Research Institute is supported by the Victorian Government’s Operational Infrastructure Support Program. We also thank Dr. rer. nat. Cédric Scherer for providing excellent resources on code-first data visualisation practices (https://www.cedricscherer.com/). This work relied heavily on the often under-appreciated work of software developers and maintainers. This work was made possible by funding from the Niels Stensen Fellowship. SnotWatch is made possible through the efforts of the SnotWatch collaboration group. The SnotWatch collaboration group includes Monash Pathology (Tony Korman), Royal Children’s Hospital Pathology (Andrew Daley, Vanessa Clifford), Alfred Pathology (Adam Jenney), Royal Melbourne Hospital Pathology (Katherine Bond), Eastern Health Pathology (Roy Chean), Northern Pathology Victoria (Yvonne Hersusianto), Barwon Health (Eugene Athan) and the Victorian Department of Health (Jim Black).

## CRediT statement

Conceptualisation: RJR, LS, KD, FC, ALP, MJ, DAL, PB, MGN, NPR, MLKT, RS, PV, TM, DPB

Data Curation: RJR, ALW, KWG, RJM, KL, LS, GM, TM

Formal Analysis: RJR, KL, GM, RJM, BN, LS, TM

Funding Acquisition: RJR, DPB, MGN, KD, ALP, FC, NPR, PB, MJ, DAL, PV, TM, JB, RS, PS

Investigation: RJR, LS, KL, ALW, JB

Methodology: RJR, LS, KL, ALW, JB, RJM, GM, KG, FC, ALP, MJ, DAL, PB, MGN, NPR, MLKT, BN, RS, PV, TM, DPB

Project Administration: RJR, LS, RJM, TM, PV, DPB

Resources: RJR, LS, KL, ALW, JB, RJM, GM, KG, KD, PS, FC, ALP, MLKT, BN, RS, PV, TM, DPB

Software: RJR, GM, TM, KL, LS

Supervision: RJR, TM, DPB, LS, PV, PB, MGN, NR, DAL, MJ, BN, RS

Validation: RJR, TM, LS, KL, FC, ALW

Visualisation: RJR, GM, TM

Writing – original draft: RJR, TM, DPB

Writing – review & editing: RJR, LS, KL, ALW, JB, RJM, GM, KG, KD, PS, FC, ALP, MJ, DAL, PB, MGN, NPR, MLKT, BN, RS, PV, TM, DPB

## Methods

### Study participants and data/sample collection

#### The Barwon Infant Study

In this study, we used samples and data from the Barwon Infant Study, a population-derived longitudinal cohort study that recruited pregnant mothers in Geelong (Victoria, Australia) between 2010 and 2013 (ethical approval project number 10/24; Barwon Health Human Ethics Committee)^22^. The data and samples used here were collected when the children were approximately four years of age. We used the subset of the cohort (inception cohort N = 1074 infants) that had both blood collection and completed questionnaires at this timepoint. The exact number of children included is reported for each separate analysis.

Body weight was measured in light clothes and without shoes using the Omron Digital Weight Scale (Model: HN-286) and recorded up to two decimal places. Height was measured by stadiometer (Seca 213 Portable Height Measuring Rod Stadiometer) and rounded to the nearest 0.1 cm. BMI was calculated as weight divided by height-squared. Body fat percentage was measured by DEXA with the Body Composition Analyzer (Tanita Bc 420 Ma). Blood samples were collected in preservative free heparinised tubes.

#### SnotWatch

SnotWatch is a collaborative platform collates results of de-identified PCR tests performed at major laboratories across the state of Victoria and has previously been described in detail^49^. SnotWatch has ethical approval for collation and analysis of de-identified PCR test results from Monash Health HREC (Reference number: RES-19-0000333L-53611). The reviewing HREC approved a waiver of consent.

For this study, PCR tests of specimens collected by Monash Health Pathology and Royal Children’s Hospital pathology between 1 December 2015 and 31 July 2018 were utilised. Test results were analysed for influenza A, influenza B, RSV, Parainfluenza 1-3, Adenovirus, human metapneumovirus and picornavirus.

#### Flow cytometry

Whole blood (heparinised; 50 µL) was stained with 3.33 µl each of CD45-PerCP (clone 2D1, BD #347464), CD4-PE (clone L200, BD #550630) and CD3-FITC (clone SK7, BD #349201) antibodies for 15 minutes at RT in the dark. Red blood cells were lysed by adding 2 mL appropriately diluted lysis solution (BD FACS Lysis solution, Cat. No. 349202) at 37 °C for 15-30 minutes in the dark. The cells were washed two times in PBS and fixed (200 µl of FIX solution (1% Formaldehyde, 2% HI-FBS, 97% PBS))) prior to acquisition on a BD FACS Canto II. Data was analysed in FlowJo v10.10.1. CD45+ events were selected and an FSC-A/SSC-A gate was used to remove spurious events and debris. Lymphocyte, monocyte, and granulocyte relative abundances were estimated based on CD45 expression in combination with SSC-A.

#### GlycA measurements

Plasma samples from samples collected in sodium heparin tubes were shipped on dry ice to Nightingale Health (Finland). GlycA was measured by NMR spectroscopy coupled with Nightingale Health proprietary algorithms. GlycA reflects the concentration and glycosylation patterns (specifically N-acetyl methyl groups) of several acute phase proteins^38^.

#### hsCRP measurements

high-sensitivity C-reactive protein (a well-known acute phase protein often regarded a benchmark measure of systemic inflammation) was measured in plasma by enzyme-linked immunosorbent assay (ELISA; R&D Systems), according to the manufacturer’s instructions.

#### Whole-blood stimulations

Within 2 hours after blood collection, an aliquot of (heparinised) whole blood was diluted 1:1 with RPMI and 180 µL was transferred to previously batch-prepared plate strips containing 20 µl of concentrated stimulus. The final concentrations and details of the used stimuli are in Table 1. The samples were incubated under standard cell culture conditions (37 °C, 5% CO2, high humidity) for 24 hours (actual range: 21h50m – 28h33m; mean 24h01m), after which they were centrifuged and the clear supernatants were stored at -80 ° C until Luminex analysis.

#### Luminex measurements and data processing

Prior to measurements, samples were randomised over Luminex plates (keeping all 8 samples of each participant together) using the Omixer package. A custom 15-plex Procartaplex (Thermo Fisher) panel was used for Luminex measurements, according to the manufacturer’s instructions. Briefly, magnetic Luminex beads were washed using a Bio-Rad Bio-Plex Pro II Wash Station and incubated with standard protein solutions or 10-fold diluted samples (in universal assay buffer) for 2 hours. The beads were washed again, and detection antibodies were added for an additional 30 minutes. After another wash, Streptavidin-PE was added to generate the reporter signal (incubation at RT for 30 minutes). Beads were resuspended in 120 µl reading buffer and acquired on a BioPlex-200 instrument.

Protein concentrations were interpolated from a standard curve using the drLumi R package. A 5-parameter log-logistic (5PLL) functions was preferred to fit the standard curve, with 4PLL serving as a backup in case of no convergence. Background handling was set to “ignore”. The built-in automatic outlier flagging was used to improve curve fit. Limits of detection (taking into account the samples were diluted 10-fold) are reported in Table 3. Cytokine concentrations were log10-transformed prior to analysis.

**Table 3:** Details of analytes measured by luminex.

| Analyte | Lower limit of detection (pg/ml) at 10x sample dilution | Upper limit of detection (pg/ml) at 10x sample dilution | Common synonyms |
| --- | --- | --- | --- |
| IFNα | 5.1 | 21000 |  |
| IFNγ | 142.3 | 583000 |  |
| IL-1β | 17.2 | 70500 | IL-1F2 |
| IL-10 | 16.4 | 67000 |  |
| IL-1Ra | 312.3 | 1279000 | IL-1RN |
| IL-5 | 68.4 | 280000 |  |
| IL-6 | 94.7 | 388000 |  |
| IL-8 | 17.8 | 73000 | CXCL-8 |
| CXCL10 | 9.0 | 37000 | IP-10 |
| MCP-1 | 35.2 | 144000 | CCL2 |
| PDGF-BB | 62.3 | 255000 |  |
| TNF | 61.5 | 252000 |  |
| VEGF-A | 43.5 | 178000 |  |

#### Quality control and outlier exclusion

We concatenated the data and employed a PCA-based method for outlier detection (implemented in factoextra) and excluded those individuals from the analysis. We furthermore excluded children with BMI > 35 (n = 1) and with excessively high neutrophil abundance indicative of active infection (n = 2), due to these conditions altering the innate immune responses. The analytes IL-37 and CCL3 were excluded from the analyses due to having almost no detectable values (nearly all measured values were too low or too high, respectively). The cytokine data is visualised in Figure 1 (heatmap and PCA) and Supplementary Figure 1.

#### Visualisation of adjusted cytokine concentrations

Adjusted cytokine concentrations for visualisation were calculated using a linear regression approach. A linear model was fit for each stimulus-cytokine combination with log10-transformed cytokine concentrations as outcome and the following as covariates (unless one of these was itself the variable of interest in the investigation): time in freezer, exact incubation time, sex, age, BMI. Adjusted values were calculated as the model residuals with the overall mean cytokine concentration added back, such that adjusted concentrations represent variation unexplained by these covariates while remaining on the original measurement scale. NB: for further clarity, statistical inference was done based on non-adjusted data using complete covariate-adjusted models.

#### Data exploration – individual cytokine plots, heatmap, and principal component analysis

To generate the individual cytokine plots in Figure S1, the cytokine data were adjusted for ‘freezer storage time of the sample’ and ‘exact incubation time of whole blood stimulation’ using the linear regression approach described above.

These adjusted concentrations were also used for generating the heatmap and PCA plots in Figure 1. For the heatmap, min-max normalisation was used to set the cytokine concentrations on a unified scale between 0 and 1. The normalised expressions were plotted on a heatmap (‘ggHeatmap’) using compete Euclidian hierarchical clustering (options: dist_method=”euclidian”, hclust_method=”ward.D2”) on both the rows (stimuli) and columns (analytes). The (not normalised) data were scaled and centered prior to PCA.

#### DNA isolation and genotyping

Genomic DNA was extracted from cord blood using the QIAamp DNA QIAcube HT Kit (QIAGEN, Germany) in line with manufacturer’s instructions. DNA was shipped on dry ice to Erasmus MC (the Netherlands) for genome-wide genotyping by the Infinium Global Screening Array-24 v1.0 BeadChip (Illumina, US).

#### Data cleaning and Imputation

Specific single nucleotide polymorphism (SNP) genotypes for each individual were imputed using the Michigan Imputation Server 2 (University of Michigan, Ann Arbor, MI, United States) using the Minimac4 imputation engine, with the Haplotype Reference Consortium (release 1.1) reference population^78^. Specific SNP genotype calls were excluded (i) pre-imputation, based on missingness (>2%), minor allele frequency (<1%), and deviation from Hardy-Weinberg equilibrium (*p* < 1e-6), (ii) post-imputation, based on missingness (>2%), imputation R-squared (<0.2), and variants that could not be confidently matched back to rsIDs. Participant samples were excluded based on missingness (>2%) and heterozygosity (>3 standard deviations (SDs) from the cohort mean)^79^.

#### Cytokine-QTL mapping

The set of SNPs was filtered for minor allele frequency > 5% (using Plink v1.9). We used the MatrixEQTL package to identify SNPs associated with cytokine production, with the following settings: the threshold for outputting results was set at *p* = 1 (as to not prune any results for downstream processing); linear models were used to determine associations; ‘pvalue.hist’ was set to ‘qqplot’ to inspect *p*-value distribution. The other settings were left at their defaults.

We removed stimulus-cytokine combinations that had > 50% of values out-of-range (representing cytokines that were not well-expressed upon exposure to that particular stimulation, and those that were expressed in extremely high concentrations relative to the other measured cytokines). Participant characteristics (exact age, sex), technical variables (exact incubation time of the whole blood stimulation, time-in-freezer of each sample), granulocyte percentage in whole blood, and circulating infection pressure (number of monthly nasal viral swabs) were used as covariates. We established three thresholds for statistical significance: p < 5 x 10^-6^ as a lenient screening threshold (justified by the relatively low number of samples and unique nature of the samples), p < 5 x 10^-8^ for genome-wide significance (a commonly used threshold), and *p* < 1.39 x 10^-9^ for study-wide significance (adjusted for the number of effective tests, as in reference^24,25^).

For Figure 2 and Supplementary Figure 2, as we were interested in identifying stimulus-specific associations, we performed cytokine-QTL mapping for each stimulus separately. Genomic location data was added to the cytokine-QTL results from the original Plink files. For the Manhattan plot, we merged the QTL-mapping results into a single dataset. We down-sampled the associations with *p* > 5 x 10^-6^ and pruned associations *p* > 0.05 (only for this visualisation) in a density-conscious manner to prevent overplotting. All associations *p* < 0.05 were plotted.

#### Variance partitioning

We used a grouped mixture of regressions software package (GMRM; developed by Orliac *et al.*^80^) to estimate the percentage of variance of cytokine responses that can be explained by genomic variants. To prevent overfitting, we used the top 50 independent SNPs as input for the model. These top independent SNPs were selected based on their *p*-value and linkage disequilibrium (LD); From SNPs that were in LD, only the SNP with the lowest *p*-value was retained (‘clumping’ as implemented in *bigsnpr::snp_clumping;* R^2^=0.2, size=500 kb). GMRM was run for 500 iterations, without using the functionality to subdivide SNPs into annotated groups. A ‘burn-in’ of 100 iterations was selected based on what was done in the original GMRM publication^80^; the final 400 iterations were used to estimate the mean and 95% confidence interval of the percent of variance explained by the top 50 independent SNPs.

#### Pathway enrichment analyses (MAGMA)

Multi-marker Analysis of GenoMic Annotation (MAGMA) was used to perform gene set enrichment analyses. First, we mapped the SNPs present in our dataset to gene locations using a window of 35 kb upstream and 10 kb downstream. For each stimulus-cytokine combination separately, we first performed the gene-level analysis step to go from ‘per-SNP’ to gene-based *p*-values. Gene set enrichment analyses were performed against the Reactome database accessed from MSigDB.

#### Establishing cytokine modules (CytoMod)

To establish cytokine modules, we used the CytoMod software^28^ made available on GitHub by Cohen *et al*., with the modifications proposed by Jack Bosco to fix compatibility issues. We used the Reticulate R package as an interface to Python from R. CytoMod offers the choice of using ‘raw’ or ‘adjusted’ cytokine concentrations for establishing the modules; we used adjusted concentrations for this manuscript. We captured the results and prepared our own versions of the output plots in R, for the purpose of streamlining figure preparation.

#### Determining associations between variables of interest and cytokine responses

We used multiple linear regression to determine associations between variables of interest and either stimulus-cytokine combinations or stimulus-specific cytokine modules. Technical covariates (exact incubation time of the whole blood stimulation in hours, and sample storage time at -80 °C) were always included in the models. Unless otherwise indicated (or when one of these was subject of the investigation itself), participant sex, exact age, and BMI were also used as covariates. We used the Benjamini-Hochberg method^81^ to correct for the number of models in each analysis, for cytokine modules. Some candidate determinants were converted to z-scores, as indicated in the text and figures.

#### Software

The following software packages were used (in no particular order):

R version 4.5.2 “[Not] Part in a Rumble”^82^ (on Windows personal computer) or R version 4.4.1 “Race for Your Life” (on HPC), with the packages tidyverse^83,84^ (core packages), haven^85^, janitor^86^, conflicted^87^, drLumi^88,89^, FactoMineR^90^, factoextra^91^, FactoInvestigate^92^, glue^93^, tidymodels^94^, rstatix^95^, corrplot^96^, khroma^97^, viridis^98^, ggheatmap^99^, gghalves^100^, ggforce^101^, ggdist^102^, ggh4x^103^, colorspace^104^, scales^105^, patchwork^106^, ggside^107^, ggrepel^108^, GGally^109^, Omixer^110^, hms^111^, readxl^112^, openxlsx2^113^, gt^114^, reticulate^115^, anthro^116^, anthroplus^117^, locuszoomr^118^, EnsDb.Hsapiens.v75^119^, BiocManager^120^, GenomeInfoDb^121^, GenomicRanges^122^, genio^123^, bigsnpr^124^, data.table^125^, MatrixEQTL^126^, foreach^127^, doMC^128^.

Other software packages used: CytoMod^28^, MAGMA^129^, PLINK^130^. We used Reactome pathway databases^131^ downloaded from MSigDB. Figure preparation was done as much as possible in R, but final compilation and non-data illustrations were prepared in Adobe Illustrator.

All code used for the analyses and figure preparation will be made available on Github or similar upon publication.

#### Statistical reporting

Primary analyses (associations between variables of interest and cytokine modules): FDR-adjusted, two-tailed *p*-values below 0.05 were considered statistically significant, except where otherwise indicated in the text. Secondary analyses (associations between variables of interest and individual stimulus-cytokine combinations): nominal two-tailed *p*-values below 0.05 were deemed significant. We acknowledge showing or reporting confidence intervals has substantial added value, but had to sacrifice this (in most cases) in the main text and in the figures in favour of clarity and readability. The regression outputs including confidence intervals of each analysis, on which the summary dotplots and volcano plots are based, are included as supplementary tables.

#### Data availability

With the approved ethics for this study, the individual participant data cannot be made freely available online. Interested parties can access the data used in this study upon reasonable request, with approval by the Barwon Infant Study data custodians. As part of this process, researchers will be required to submit a project concept for approval, to ensure the data is being used responsibly, ethically, and for scientifically sound projects.

## Supplementary figure captions

**Supplementary Figure 1: all measured cytokine responses.** The graphs are ordered by stimuli (columns) and analytes (rows). The depicted values were adjusted for ‘time in freezer’ and ‘exact incubation time’ using a linear regression approach (see Methods section).

**Supplementary Figure 2: Genetic variants and cytokine responses.** (A) Locuszoom plot (top) and genetrack (bottom) of rs72679554. (B-C) Covariate-adjusted concentrations of respectively IL-10 and TNF upon stimulation with Lyovec/dsRNA, in children with different genotypes on this location. (D) Locuszoom plot (top) and genetrack (bottom) of rs2384950. (E) Covariate-adjusted concentrations of IFNγ upon stimulation with R848, in children with different genotypes on this location. (F) Locuszoom plot (top) and genetrack (bottom) of rs2300875. (G) Covariate-adjusted concentrations of MCP1 upon stimulation with poly(I:C), in children with different genotypes on this location. (H) Locuszoom plot (top) and genetrack (bottom) of rs6112096. (I) Covariate-adjusted concentrations of MCP1 upon stimulation with RPMI alone, in children with different genotypes on this location. (J) Top 10 Reactome pathways that were enriched (nominal *p* < 0.05), shared between the highest proportion of cytokines for stimulation with each of respectively PGN, R848, poly(I:C), Lyovec/RPMI, and Lyovec/dsRNA. In this figure, covariates included in the models are time in freezer, exact incubation time, sex, seasonal infection burden, and granulocyte percentage. Boxplots are in the style of Tukey.

**Supplementary Figure 3: Correlation between individual cytokines and module scores** Each panel represents a cytokine module, organised by stimulus and from left to right. The stimulus and module are indicated in the lower-right subpanel on the diagonals. The Pearson correlation coefficients are indicated by the numbers in top subpanels, while the individual data points (log-transformed and standardised) are plotted in the lower subpanels.

**Supplementary Figure 4: Associations between birth variables and cytokine responses.** (A) Volcano plot indicating the change in cytokine levels with an increase of 100 gram in birthweight. (B) Volcano plot indicating the change in cytokine levels with an increase of 1 week gestational age. (C) Volcano plot indicating the change in cytokine levels with an increase of 1 SD in birthweight-by-gestational-age. (D) Dotplot summarising regression results between each variable of interest and cytokine module expression score (reference: vaginal birth, except the comparison between ‘with’ vs ‘no’ labour in Caesarean birth). ‘Effect size’ was calculated as absolute value of the regression estimate. In this figure, ‘covariate adjusted’ includes time in freezer, exact incubation time, and sex, age, and BMI.

**Supplementary Figure 5: Associations between systemic inflammation/cell type composition and cytokine responses.** (A-C) Distribution of cell type percentage in whole blood for respectively granulocytes, lymphocytes, and monocytes. (D) Comparative plot indicating the difference change in module expression scores when GlycA or hsCRP is increased by 1 SD. The points are coloured by the indicated FDR-adjusted *p*-value significance categories. (E) Comparative plot indicating the difference change in cytokine levels when GlycA or GLR is increased with 1 SD. The points are coloured by the indicated FDR-adjusted *p*-value significance categories. For panels D-E, points in the white areas are concordant between measures; points in the grey areas are discordant. (F) Dotplot summarising regression results between each variable of interest and cytokine module expression score. In each dot ‘*’ indicates FDR-adjusted *p* < 0.05. ‘Effect size’ was calculated as absolute value of the regression estimate. (G-I) Volcano plots indicating the change in individual cytokine levels upon an increase of 1 SD in respectively granulocyte, lymphocyte, or monocyte percentage. Panels G-I have nominal *p*-values. In this figure, ‘covariate adjusted’ includes time in freezer, exact incubation time, and sex, age, and BMI. Boxplots are in the tradition of Tukey.

**Supplementary Figure 6: Seasonal variation in cytokine module expression scores.** The graphs are organised by stimulus (rows) and cytokine modules (columns). There are empty spots in case there were fewer than 5 modules for a stimulus. The depicted values were adjusted for ‘time in freezer’, ‘exact incubation time’, sex, age, and BMI using a linear regression approach (see Methods section).

**Supplementary Figure 7: Seasonal variation in individual cytokines across stimuli.** The graphs are organised by stimulus (columns) and cytokines (rows). The depicted values were adjusted for ‘time in freezer’, ‘exact incubation time’, sex, age, and BMI using a linear regression approach (see Methods section).

**Supplementary Figure 8: Seasonal infections in the community**. (A) Number of positive viral swabs over the study period. (B) Number of total viral swabs over the study period. (C) Percentage of viral swabs that was positive over the study period.

