## Supplementary figures and images for "Genetic, intrinsic, and environmental determinants of innate immune cytokine responses in healthy four-year-old children"

### Figure 1.tif

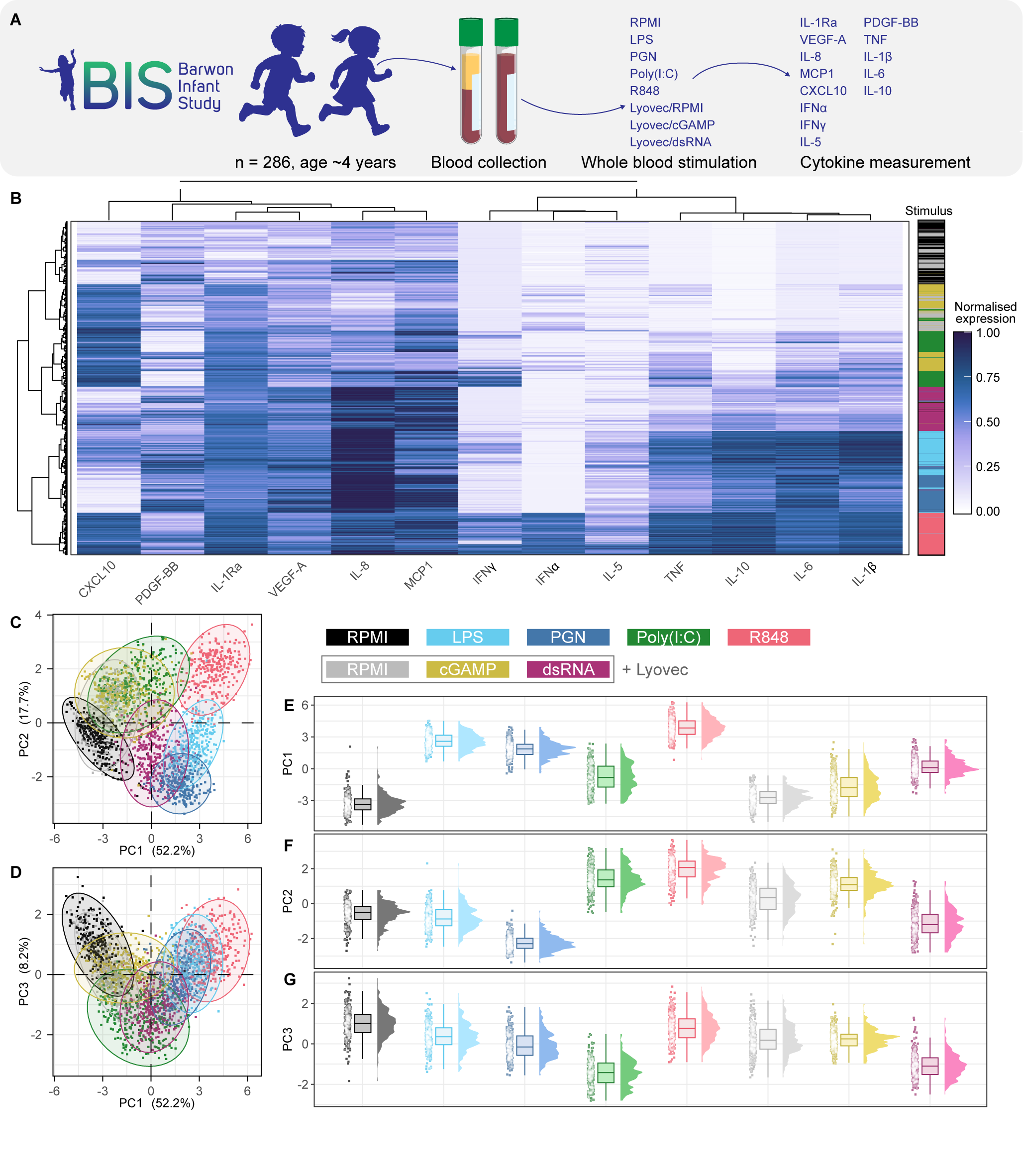

### Figure 2.tif

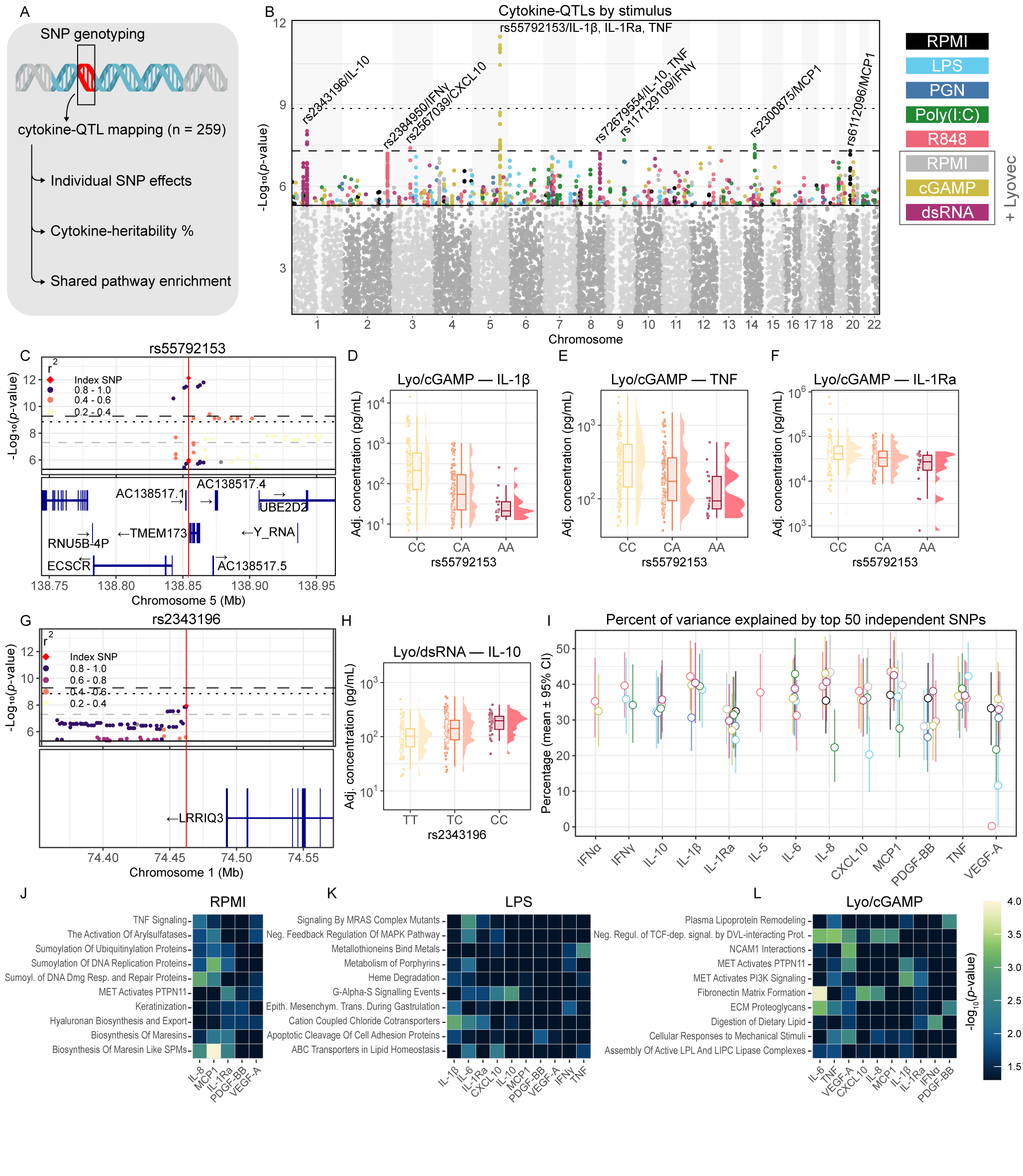

### Figure 3.tif

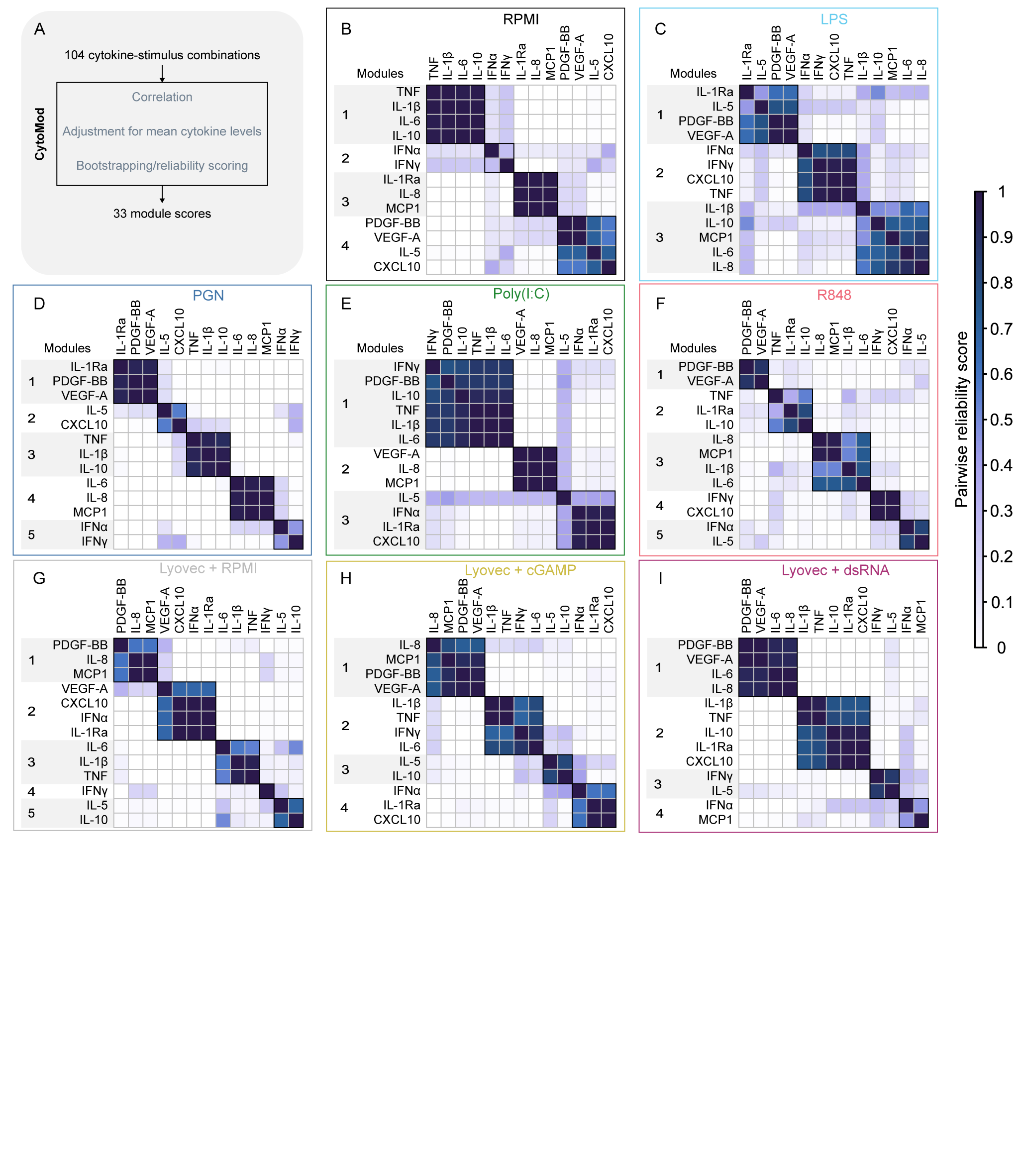

### Figure 4.tif

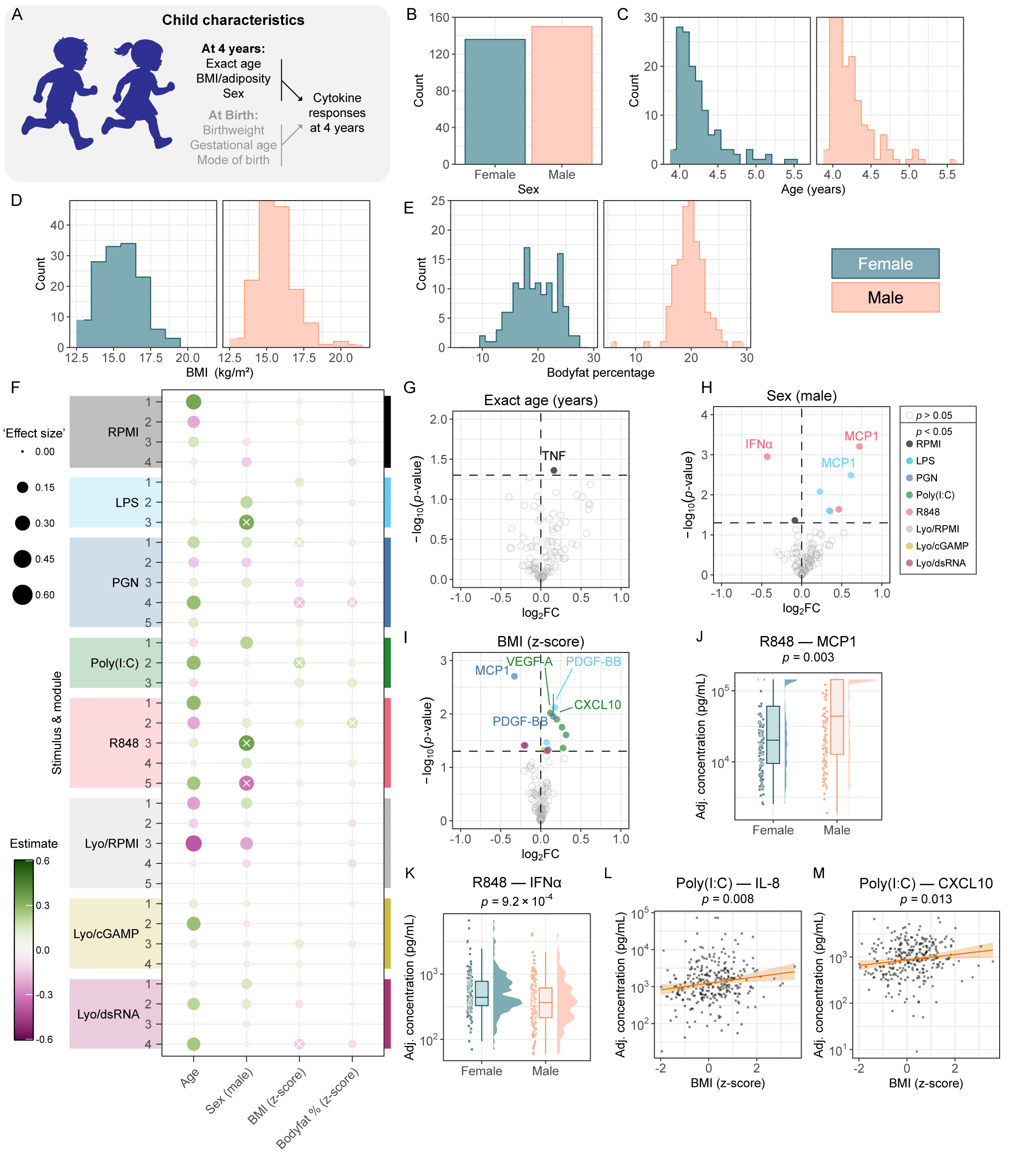

### Figure 5.tif

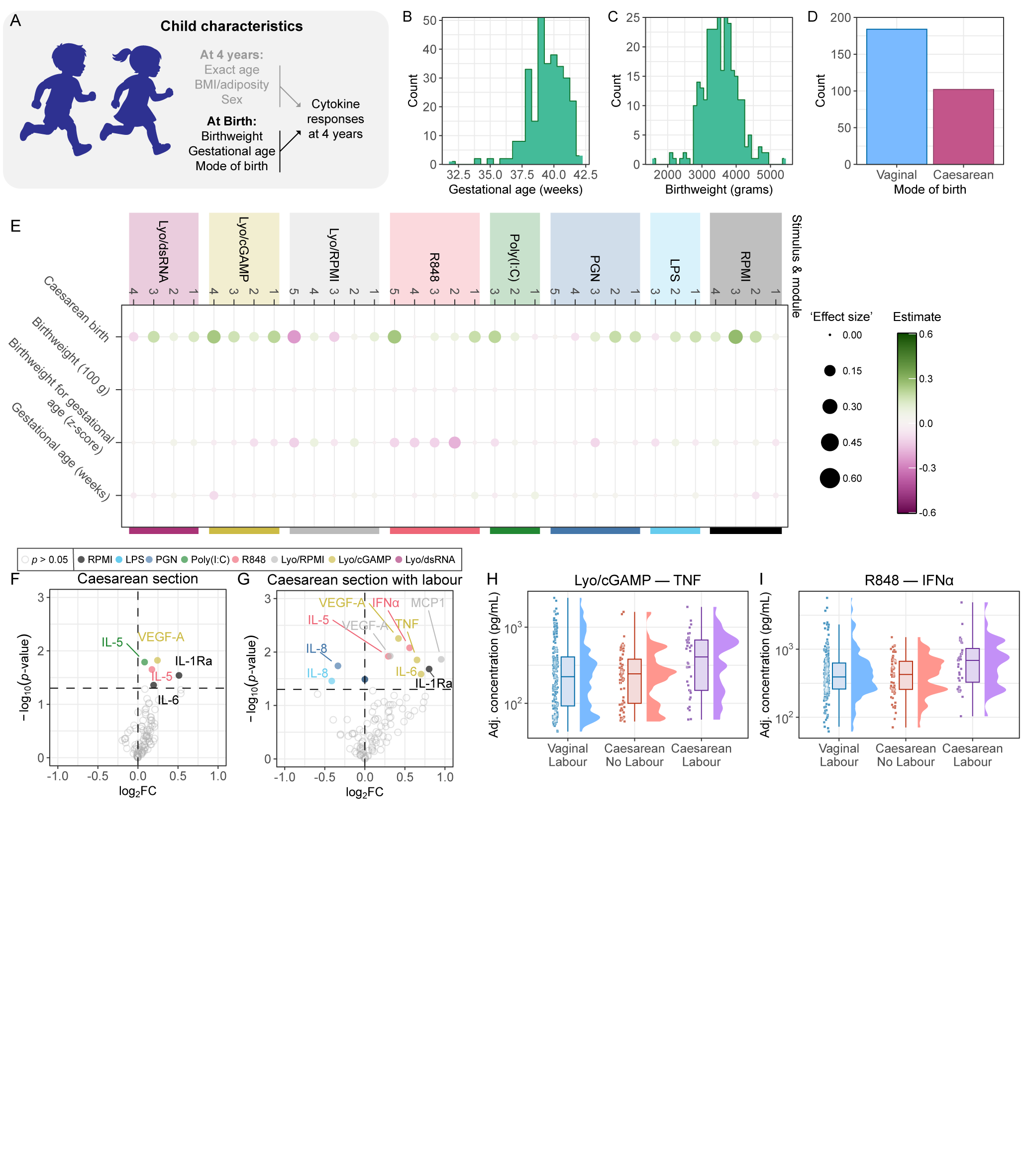

### Figure 6.tif

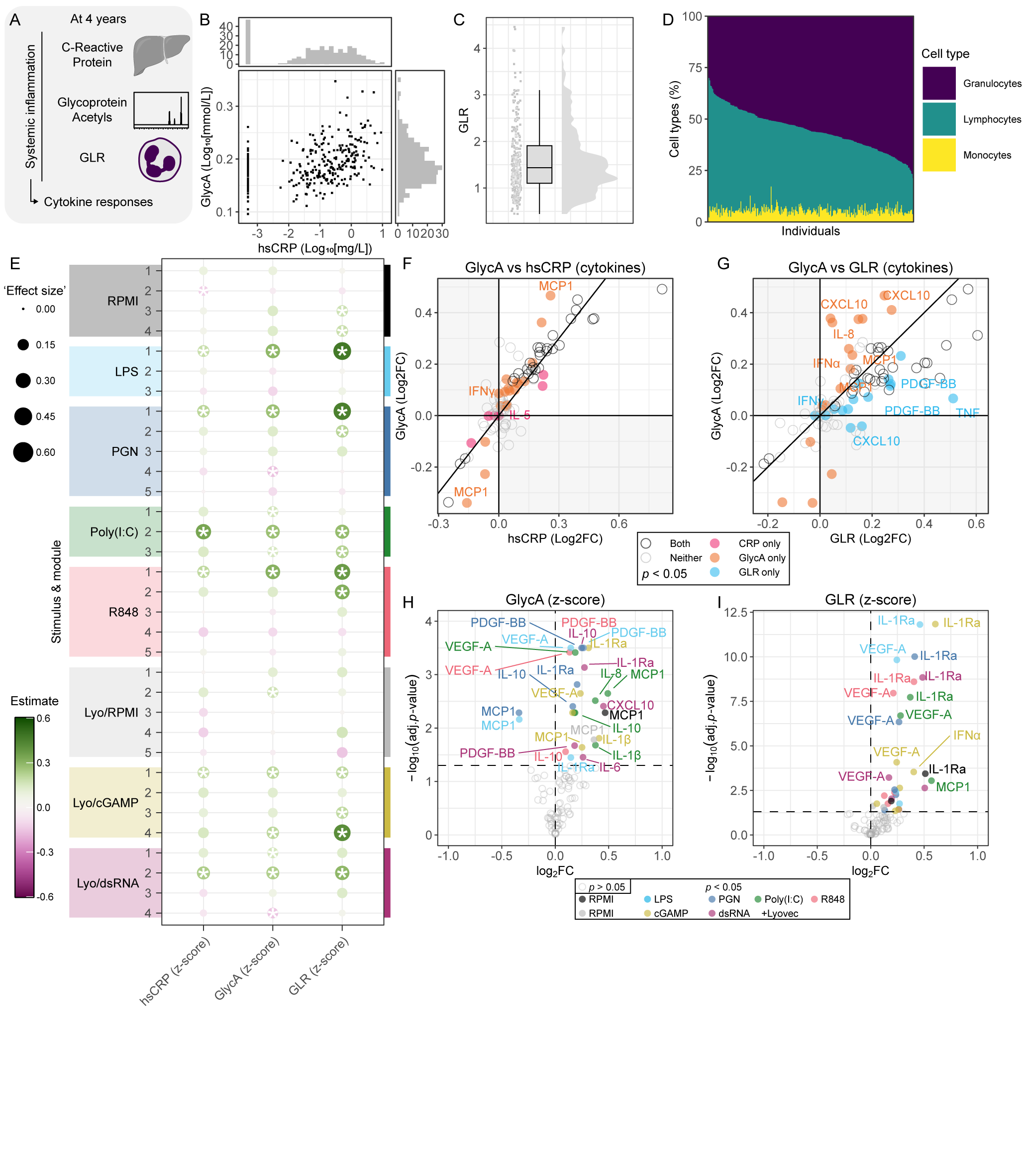

### Figure 7.tif

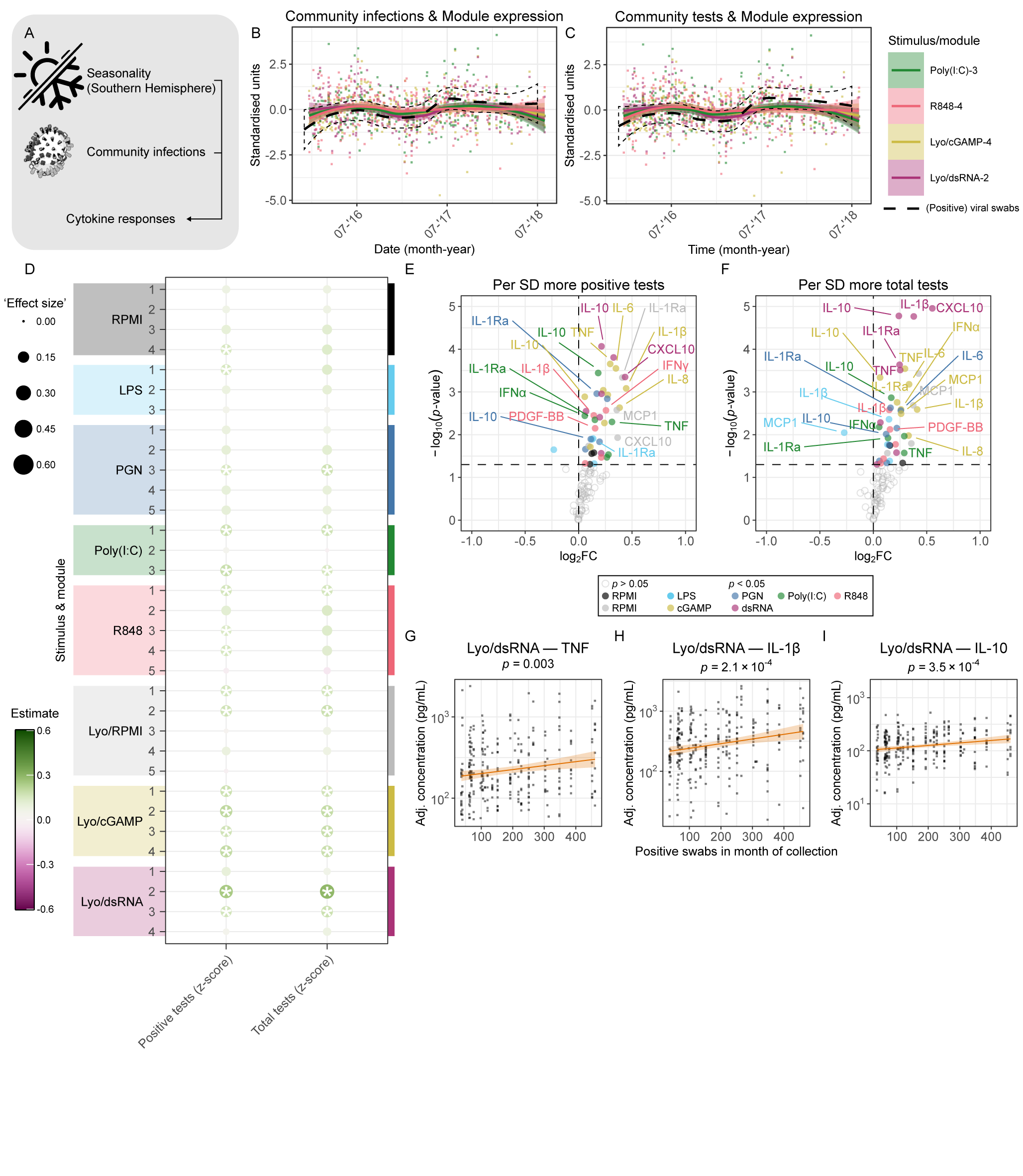

### Figure S1.pdf

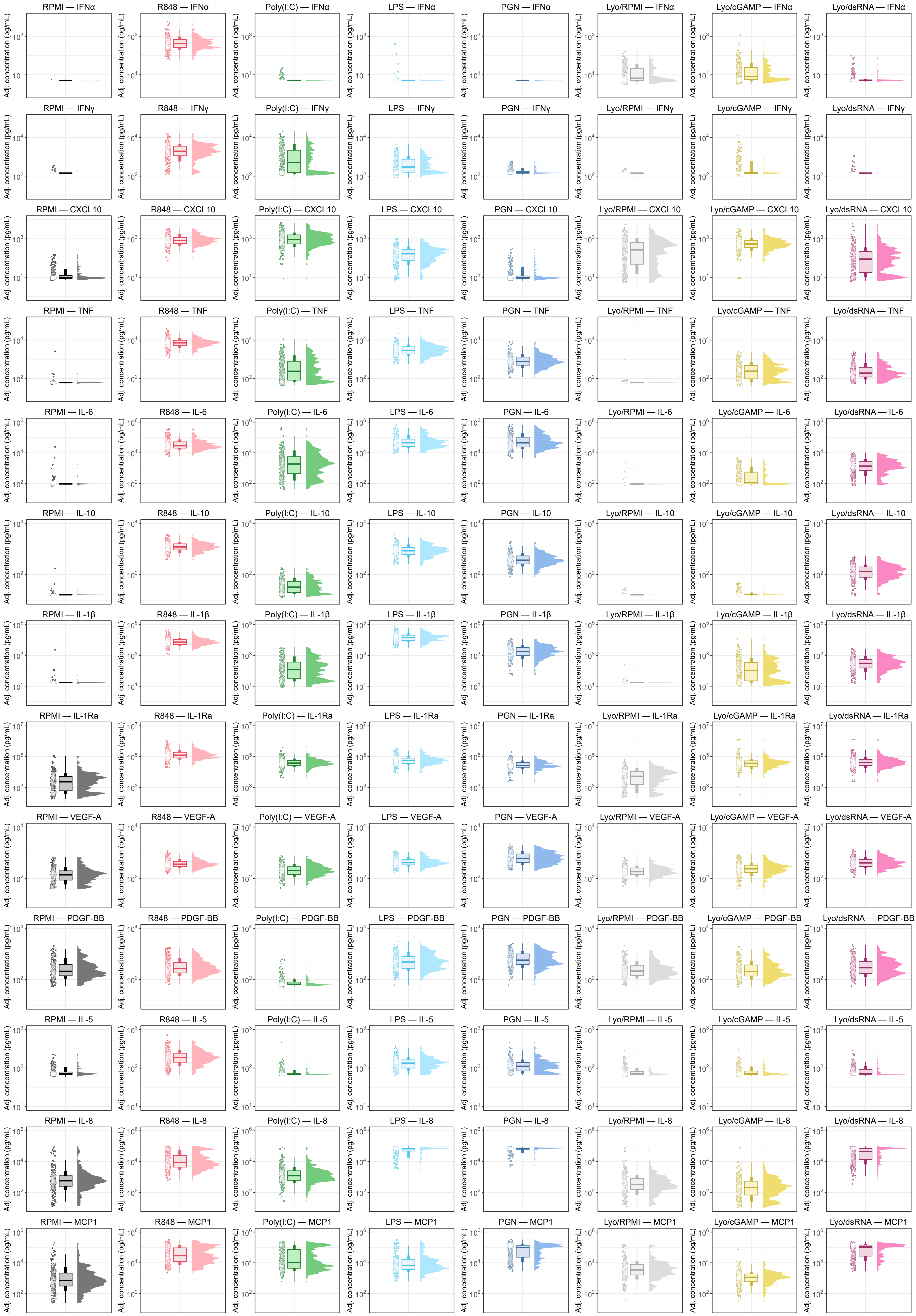

### Figure S2.tif

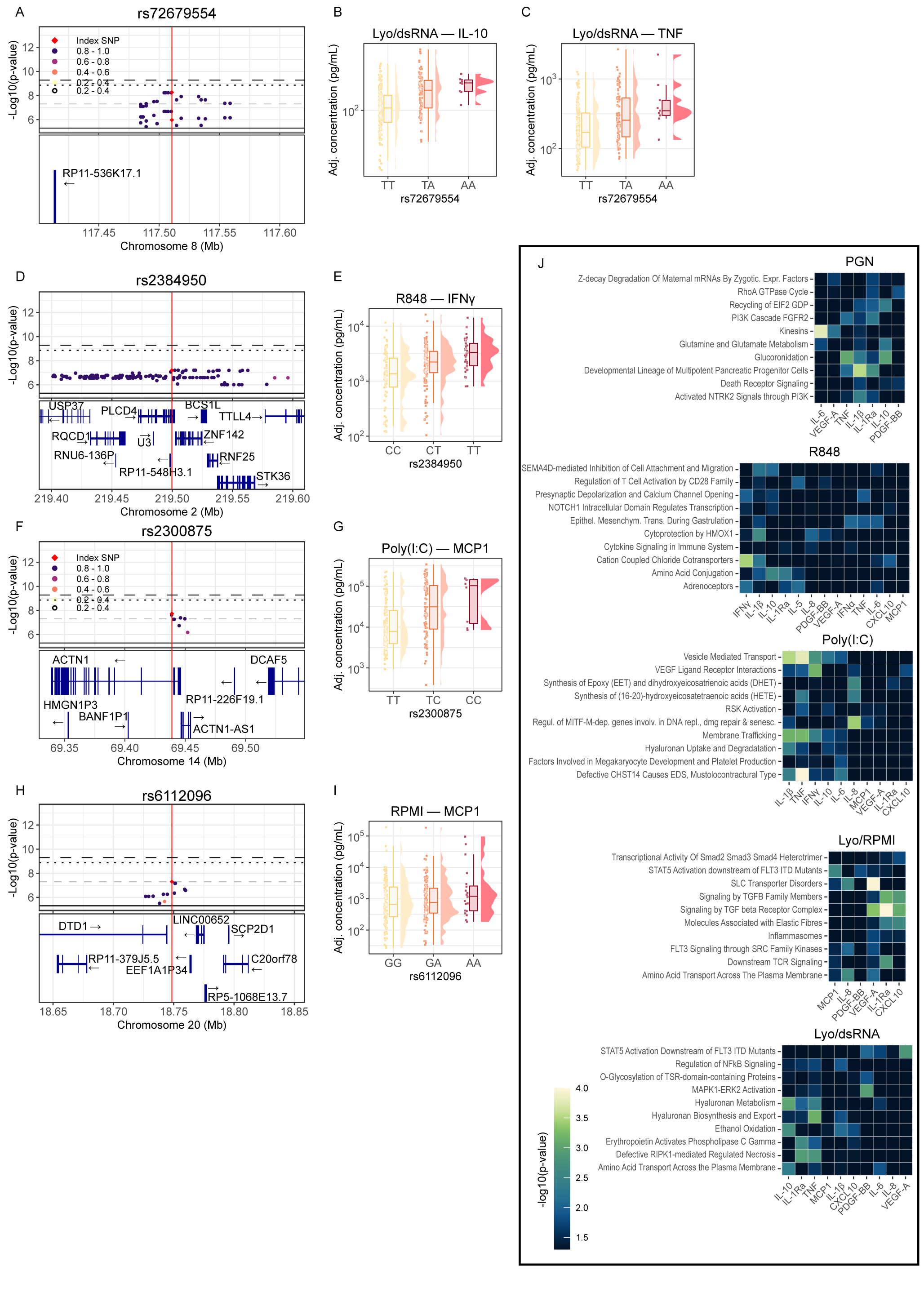

### Figure S3.jpg

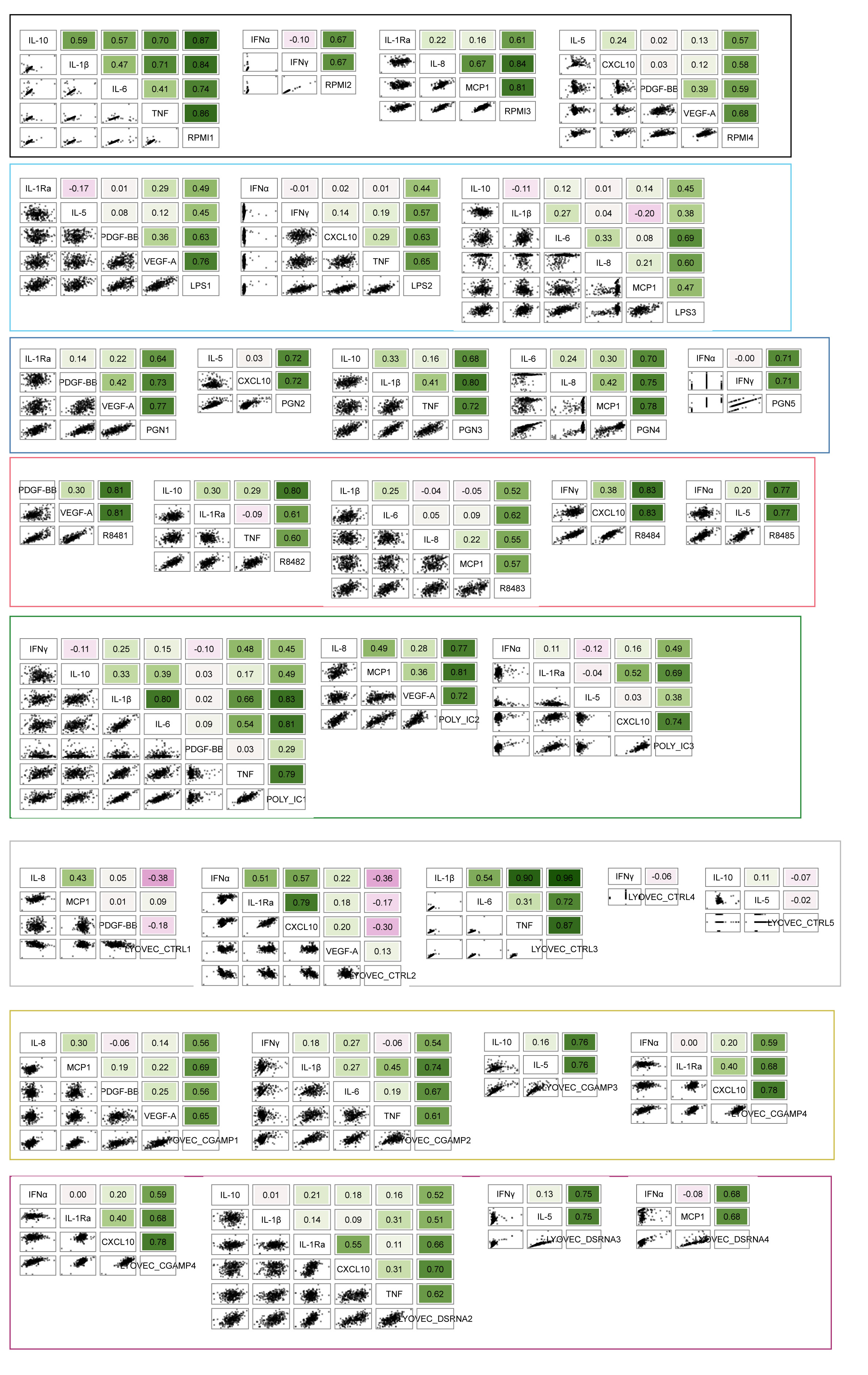

### Figure S4.tif

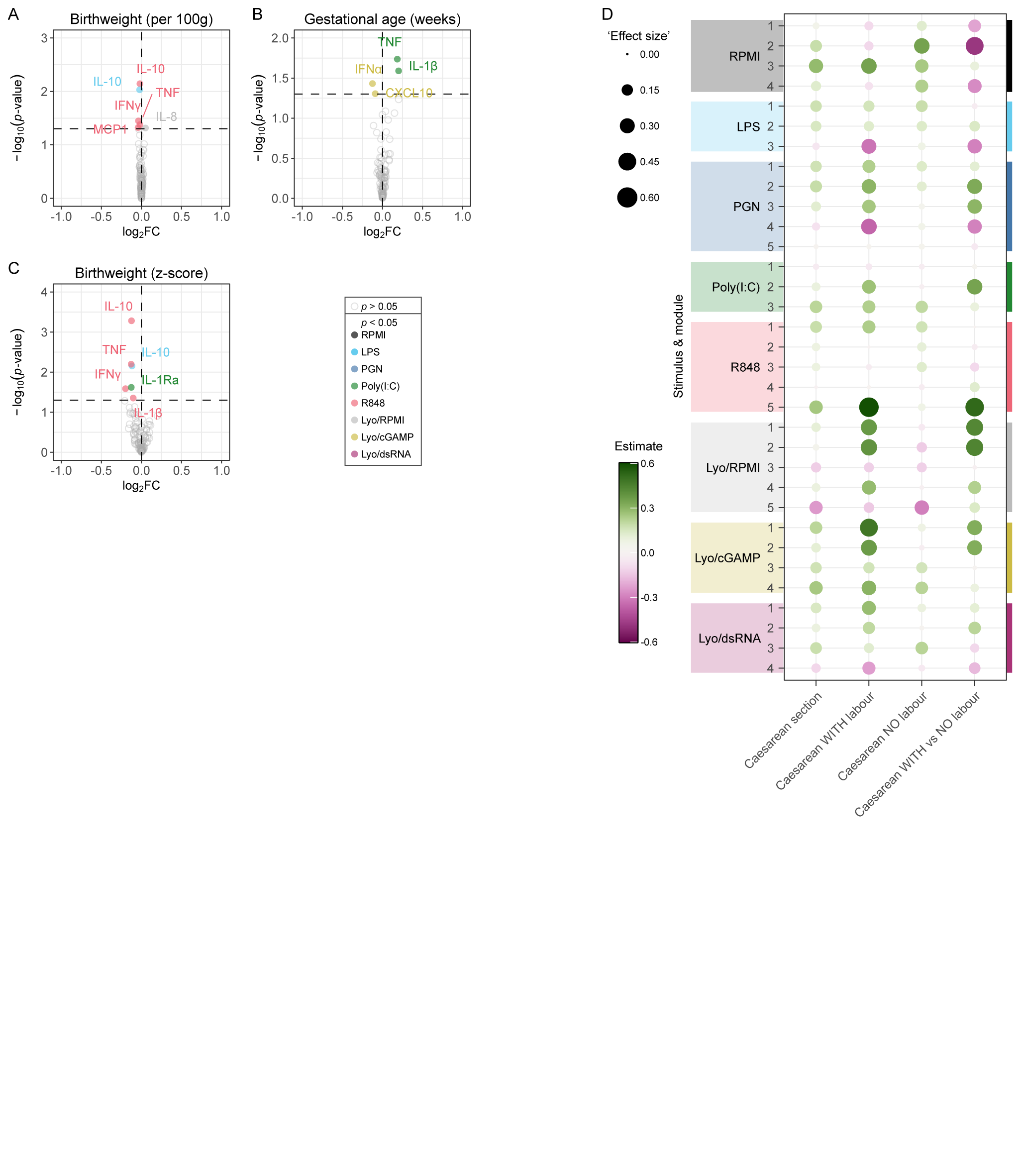

### Figure S5.tif

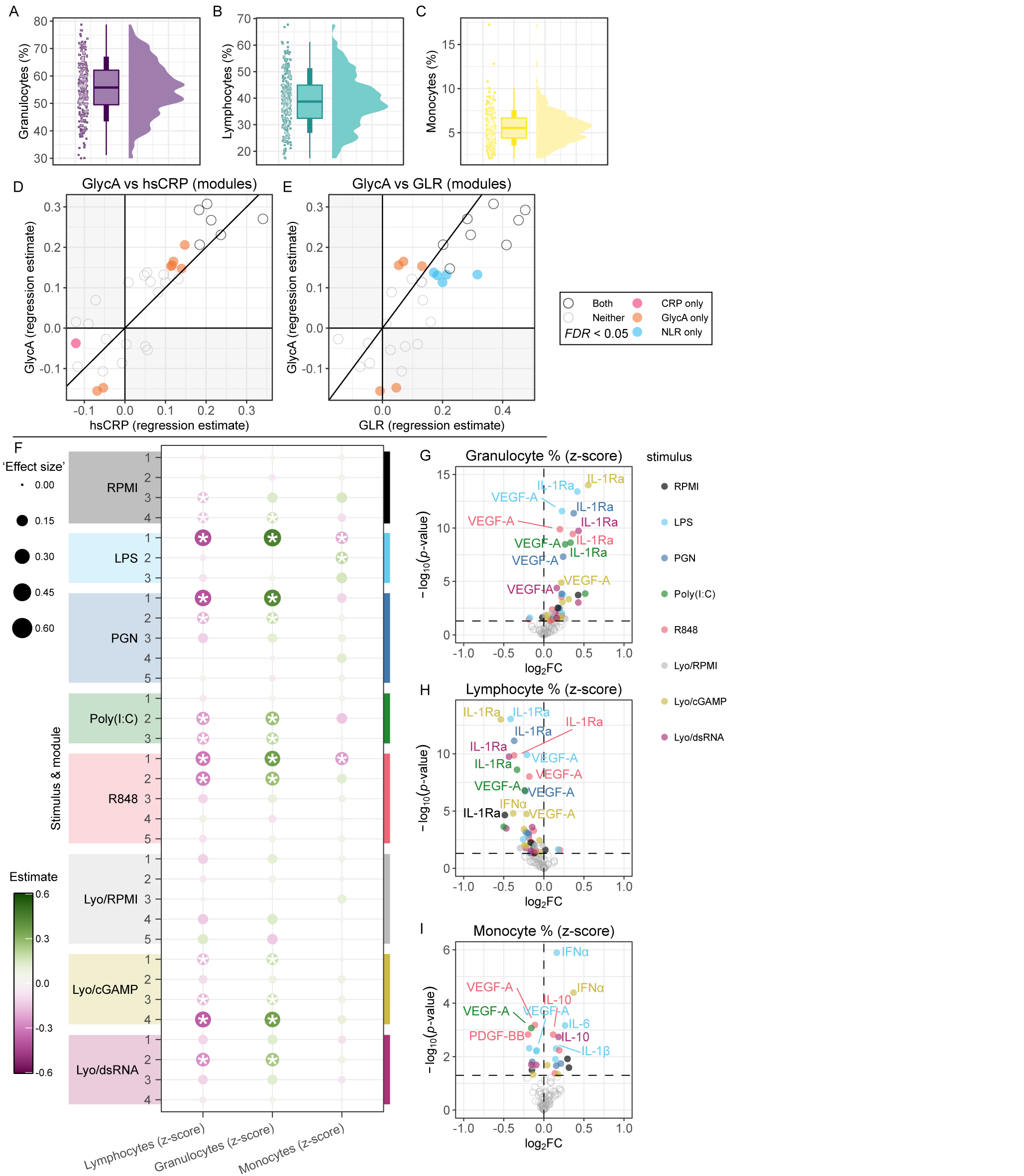

### Figure S6.jpg

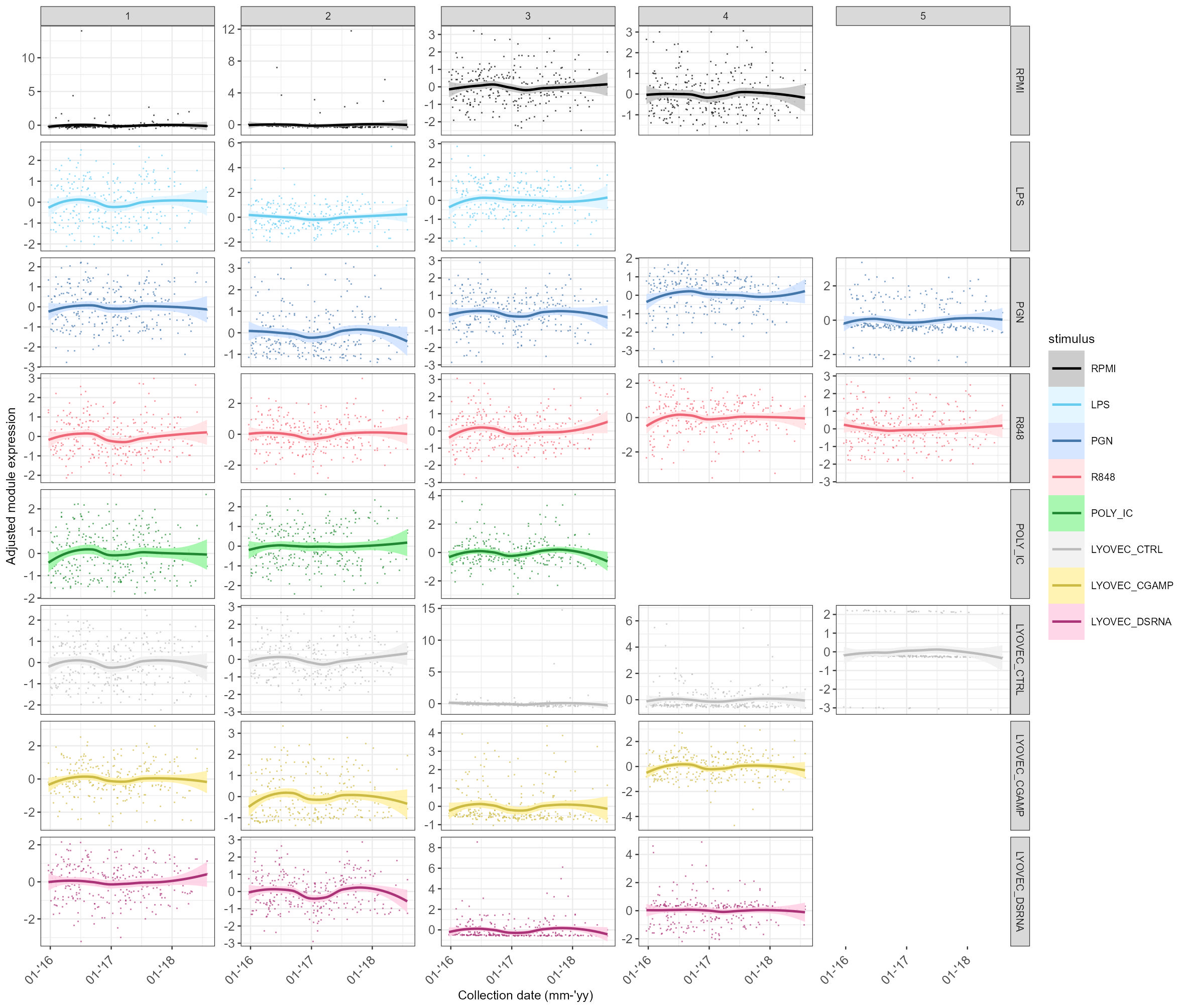

### Figure S7.jpg

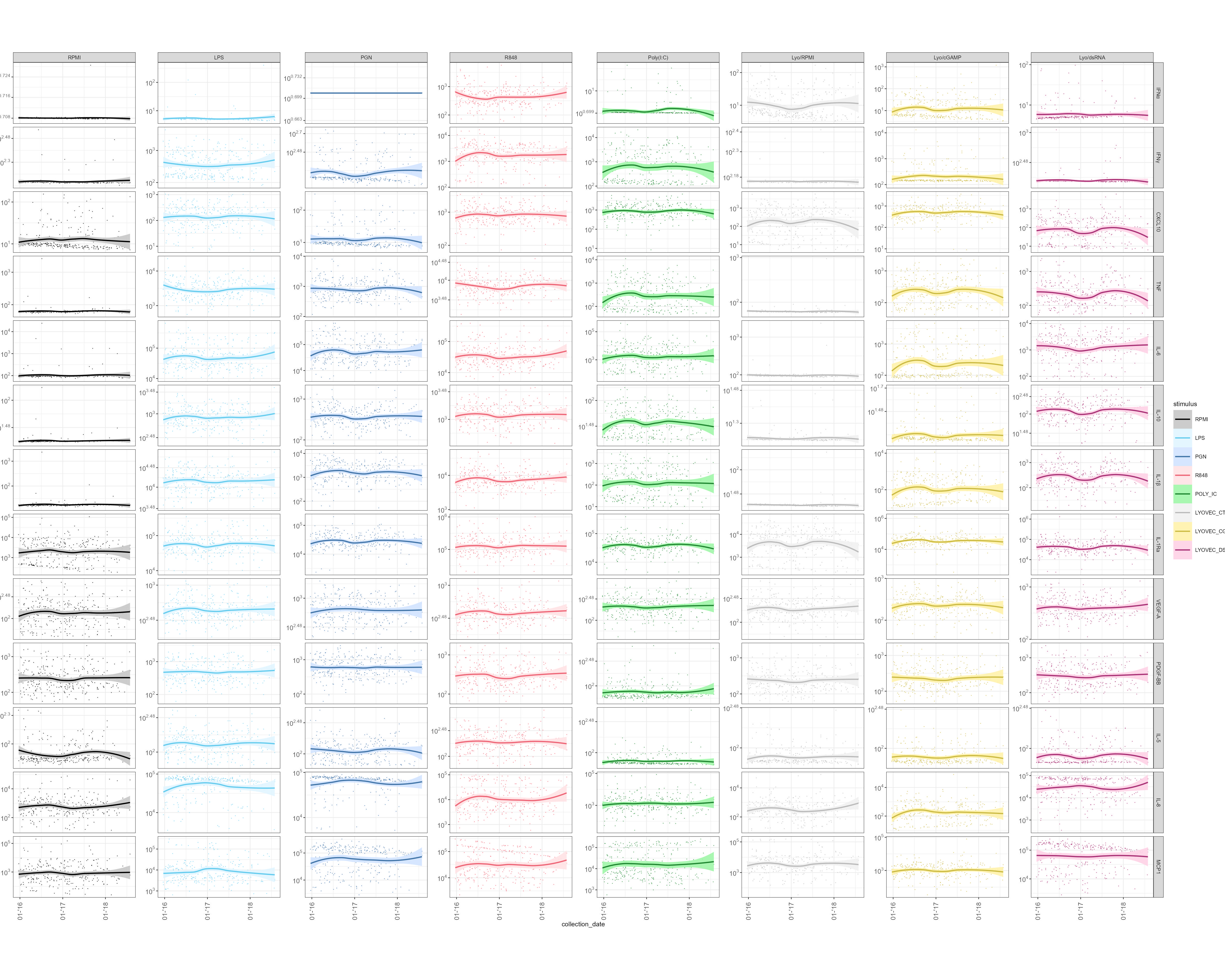

### Supplementary Figure 8.tif

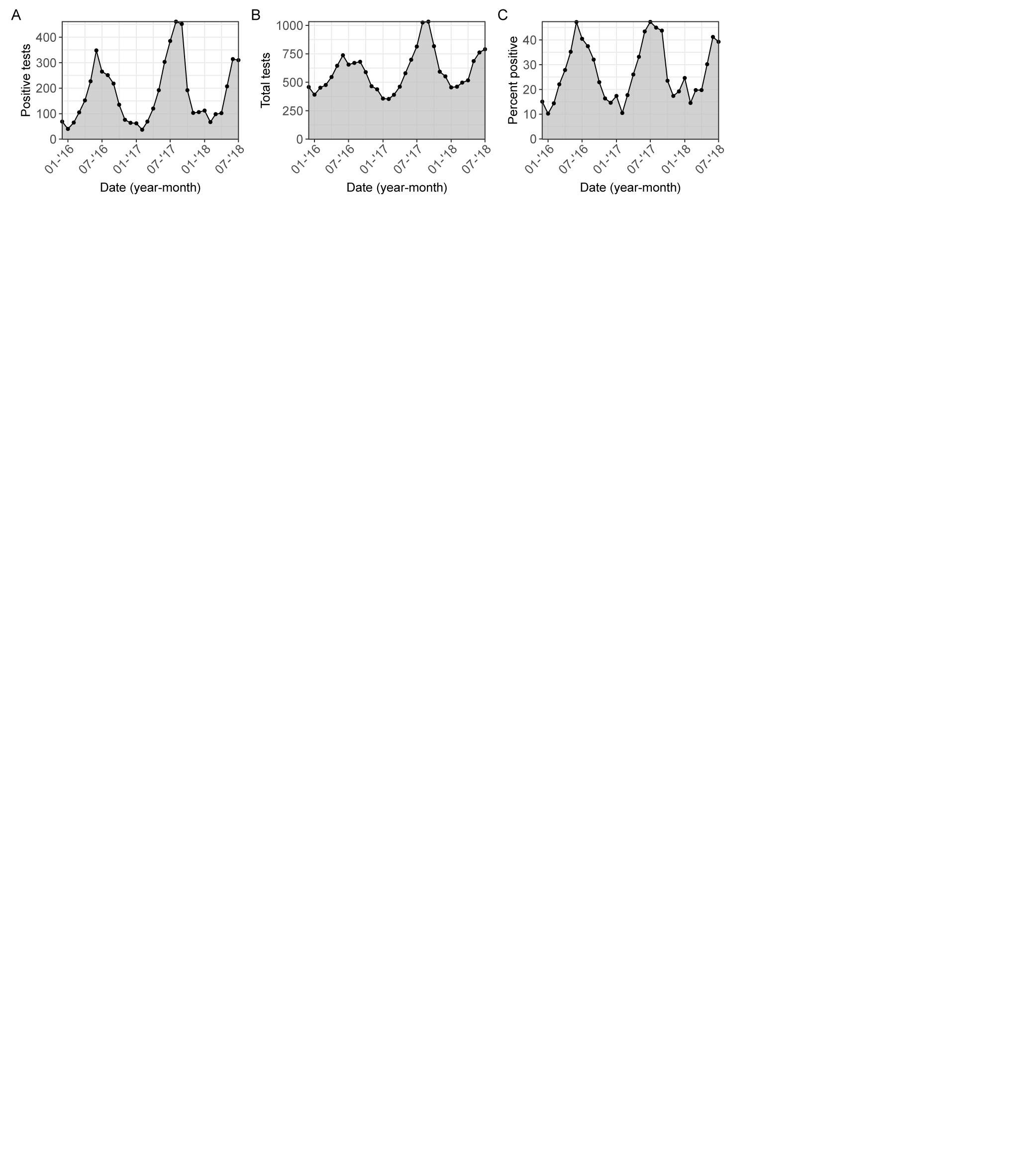
